# FUSED: A Functional Representation for Joint Structural and Elemental Analysis of Protein Ligand Binding Sites

**DOI:** 10.64898/2026.07.10.737806

**Authors:** T.M. Sajith Priyankara, Leif Ellingson

## Abstract

Ligand binding site representations are central to the analysis of protein–ligand interactions, with applications in functional characterization, binding-site comparison, and ligand recognition. Many descriptor-based approaches characterize ligand binding sites using a fixed distance threshold from the ligand, despite substantial variability in how such thresholds are defined and the possibility that relevant structural and compositional information changes across spatial scales.

We propose Functional Unification of Structural and Elemental Descriptors (FUSED), a multivariate functional representation that jointly models structural and elemental compositional information of ligand binding sites as functions of distance from the ligand. Structural information is captured through covariance-based descriptors derived from the CDPA framework, while chemical composition is represented through isometric log-ratio coordinates to account appropriately for compositional geometry. Treating distance from the ligand as a functional domain allows structural and compositional characteristics to be examined across distance thresholds rather than at a single prespecified value. The resulting representation can be used directly or combined with dimension-reduction, statistical-learning, and/or machine-learning procedures, with the distance interval tailored to the dataset, analytical task, and procedure.

We evaluate FUSED on three benchmark datasets spanning complementary ligand binding-site analysis tasks: the Extended Kahraman dataset for multiclass ligand discrimination, TOUGH-C1 for binary binding-site classification, and TOUGH-M1 for pairwise matching of pockets associated with drug-like ligands. FUSED supports strong discrimination in the EK and TOUGH-C1 tasks using standard statistical-learning procedures, while a supervised Siamese neural network applied to the full FUSED representation achieves a mean ROC-AUC of 0.9375 on TOUGH-M1 under repeated sequence-cluster-disjoint evaluation, approaching the strongest reported benchmark performance. These results demonstrate that FUSED provides a flexible, alignment-free representation that supports both direct examination of threshold-dependent binding-site characteristics and competitive downstream classification and pocket matching while remaining computationally practical.

## 1. Introduction

Computational analysis of ligand binding sites (LBSs) plays a central role in structural biology, chemical informatics, and drug discovery, as the local environment surrounding a bound ligand often contains critical information about molecular recognition, binding specificity, and biological function. A ligand binding site is commonly defined as the collection of protein atoms located within a specified distance of a bound ligand, representing the local structural and chemical environment relevant to intermolecular interactions [34, 41]. However, this seemingly simple definition introduces an important methodological challenge: the representation of an LBS depends fundamentally on the chosen distance threshold.

Distance thresholds used to define LBSs vary substantially across the literature. Thresholds near 4.5 Å have been used in several studies to characterize local binding environments [7, 48], while other work has suggested that broader neighborhoods exceeding 10 Å may be necessary to capture biologically meaningful binding-site context, particularly in druggability studies [29]. For the extended Kahraman (EK) dataset, Hoffmann et al. identified 5.3 Å as an effective threshold for classification using their structural comparison framework [22], and this convention has been adopted in subsequent studies [10, 41]. These differing choices reflect a broader conceptual issue: a fixed threshold may either exclude informative structural or chemical context or incorporate additional information whose relevance may depend on the system and analytical objective. More fundamentally, both structural geometry and chemical composition change as the neighborhood around the ligand expands, suggesting that a single-threshold representation may provide only a partial view of the binding environment.

A wide range of computational approaches have been developed for analyzing LBSs. Existing methods include shape-based approaches that characterize surface geometry [31, 39, 52], structural alignment and pocket-matching methods that compare binding sites through pairwise geometric correspondence [5, 22, 33, 53], graph-based representations that encode structural relationships among residues or atoms [38, 40, 51], and more recent machine learning approaches, including deep neural architectures trained on structural representations of protein-ligand systems [12, 25, 28]. These approaches have produced strong predictive performance in many settings, but different classes of methods involve different practical tradeoffs. For example, structural alignment and pocket-matching approaches may require computationally intensive pairwise comparisons, while more complex machine-learning approaches may rely on task-specific architectures and learned representations that can be more difficult to examine directly.

Among alignment-free approaches, the Covariance of Distances to Principal Axes (CDPA) representation provides a computationally efficient alternative by summarizing ligand binding site geometry through covariance-based structural descriptors [41]. CDPA avoids pairwise structural alignment and provides an explicitly defined geometric representation, but important limitations remain. First, CDPA focuses exclusively on structural dispersion and does not explicitly account for chemical composition, despite the well-established importance of local chemistry in ligand recognition. Incorporating such information also requires appropriate treatment of compositional geometry, since elemental proportions reside naturally in a simplex space rather than an ordinary Euclidean space. Second, CDPA is constructed at a fixed threshold and therefore does not capture how structural or compositional characteristics evolve as progressively more distant atoms are incorporated.

Several existing approaches incorporate ligand binding site information across multiple spatial scales, for example, through distance-binned physicochemical descriptors or voxel-based spatial representations [1, 26, 30, 35, 42]. However, these approaches generally encode multiscale information through predetermined spatial summaries rather than treating distance from the ligand as a functional domain over which structural and compositional descriptors are jointly represented.

In this work, we introduce FUSED (**F**unctional **U**nification of **S**tructural and **E**lemental **D**escriptors), a ligand binding site representation that jointly models structural and compositional information across a continuum of distance thresholds. Structural information is represented through CDPA-derived descriptors, while chemical composition is incorporated using isometric log-ratio (ILR) transformed compositional summaries that respect the underlying simplex geometry. Rather than representing each binding site only at a single distance threshold, FUSED treats distance from the ligand as a functional domain, allowing both descriptor families to be represented as threshold-indexed functions. This framework enables direct examination of how structural and compositional characteristics evolve across distance while retaining the flexibility to focus subsequent analyses on portions of the distance domain that are informative for a particular dataset, analytical objective, or procedure.

This functional perspective provides several advantages. It allows complementary structural and compositional signals to be analyzed jointly, allows threshold-dependent binding-site characteristics to be examined directly, and yields an alignment-free representation compatible with a range of downstream statistical and machine-learning procedures. The distance interval used for a particular analysis can be tailored to the dataset, analytical objective, and procedure rather than assumed to be universally optimal. We evaluate the proposed framework across three complementary benchmark settings: multiclass ligand discrimination using the EK dataset, binary binding-site classification using TOUGH-C1, and pairwise pocket matching using TOUGH-M1, which is restricted to pockets associated with drug-like ligands. Across these tasks, FUSED supports competitive classification performance using standard statistical-learning procedures and strong supervised pocket-matching performance when the full representation is used with a Siamese neural network, while remaining alignment-free and computationally practical. To our knowledge, covariance-based ligand binding site representations that jointly model structural and compositional descriptors as functions over distance thresholds have not previously been explored.

The remainder of the paper is organized as follows. Section 2 describes binding-site construction, the benchmark datasets, and the associated preprocessing steps. Section 3 introduces the structural and compositional descriptors used to construct the representation, while Section 4 evaluates the adequacy of their joint use at fixed distance thresholds. Section 5 defines the FUSED representation and describes its functional implementation. Section 6 illustrates exploratory and descriptive analyses of the resulting functions, including multivariate functional principal component analysis, and Section 7 examines how the choice of distance interval can affect subsequent analyses. Section 8 evaluates FUSED for multiclass and binary binding-site classification using the EK and TOUGH-C1 datasets, respectively. Section 9 considers the more application-oriented TOUGH-M1 pairwise pocket-matching task, including both direct similarity based on the FUSED representation and supervised learning with a Siamese neural network. Section 10 examines computational considerations, and Section 11 discusses the broader implications, limitations, and directions for future work.

## 2 Binding site construction and data

This section formalizes ligand binding sites as a function of the distance threshold and describes the datasets used in our analysis. We begin with the definition of the binding site in terms of protein atoms within a specified distance of the ligand, followed by descriptions of the Extended Kahraman, TOUGH-C1, and TOUGH-M1 datasets, along with the primary uses of each for this study.

### 2.1 Binding sites as a function of the distance threshold

An LBS is defined as all atoms in the protein within a specified distance threshold of the atoms in the ligand molecule. To formalize this, let *n*_*l*_ and *n*_*p*_ denote the number of atoms in the ligand and in the protein chain associated with the ligand. Let

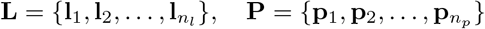

denote the sets of 3D coordinates of the ligand and protein atoms, where **l**_*j*_, **p**_*i*_ ∈ ℝ^3^. The Euclidean distance between ligand atom **l**_*j*_ ∈ **L** and protein atom **p**_*i*_ ∈ **P** is given by

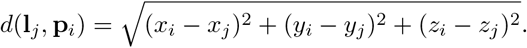

We then define the ligand binding site at threshold *t*, denoted **P**_**L**_(*t*), as the set of all protein atoms {**p**_*i*_} such that

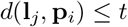

for at least one ligand atom **l**_*j*_ ∈ **L**. For consistency across all proteins, we ignore hydrogen atoms, as they are often not detected in protein crystallography [11].

For any *t*_1_ *< t*_2_, we have **P**_**L**_(*t*_1_) ⊆ **P**_**L**_(*t*_2_). Thus, as *t* increases, the binding site expands, and all information present at smaller thresholds is retained at larger ones. However, atoms included at larger values of *t* are necessarily farther from the ligand and may provide different amounts or types of information depending on the binding site and analytical objective. As a result, increasing *t* incorporates additional structural and compositional context whose usefulness may vary across analyses. This creates a fundamental trade-off between capturing relevant structural information and incorporating extraneous detail when defining binding sites using a fixed threshold.

### 2.2 Extended Kahraman dataset

The Extended Kahraman (EK) dataset was introduced in [22] as an extension of a smaller set first described in [27]. All protein structures in this dataset were obtained from the Protein Data Bank (PDB) [2, 21] using X-ray crystallography. The original dataset consists of 965 ligand binding sites (LBSs) associated with the ligand groups summarized in Table 1. The STRD group consists of LBSs that bind to either the EST or AND ligands, which are combined into a single category due to their similarity and limited sample sizes [10, 22, 41].

**Table 1:** Composition of the EK dataset before and after preprocessing.

| Ligand | AMP | ATP | FAD | FMN | HEM | NAD | STRD | GLC | PO4 |
| --- | --- | --- | --- | --- | --- | --- | --- | --- | --- |
| Original dataset | 63 | 78 | 79 | 58 | 113 | 91 | 6 | 88 | 389 |
| Analyzed dataset | 60 | 78 | 79 | 57 | 112 | 90 | — | 77 | 351 |

The ligands in this dataset vary substantially in both size and flexibility. For example, PO_4_ is small and highly rigid, while FAD is considerably larger and more flexible. This range of characteristics leads to corresponding variation in the structural and compositional properties of the associated binding sites, making the dataset well-suited for examining how binding site representations capture patterns both within ligand groups and across different ligand types. We therefore use the EK dataset to illustrate the proposed FUSED representation in detail, including visualizing group-level behavior, variability across the threshold domain, and dimension reduction. We also use it to compare the discriminatory performance of simpler single-threshold structural and compositional representations with the richer functional representation in a multiclass classification setting, providing a controlled setting for assessing what is gained as the representation becomes more comprehensive.

For the present study, we excluded the steroid (STRD) group due to its very limited sample size, leaving 959 candidate LBSs. Since the proposed FUSED representation requires both structural and compositional descriptors to be well-defined over the functional domain, additional preprocessing was required. A lower threshold of 4.8 Å was selected because smaller thresholds resulted in some binding sites containing too few atoms for stable descriptor construction. We further excluded binding sites with fewer than 10 atoms at this threshold, as reliable CDPA estimation requires sufficient structural observations. Because the ILR compositional descriptors require strictly positive proportions for carbon, oxygen, and nitrogen, binding sites with zero counts in any of these components at 4.8 Å were also removed. After filtering, 904 binding sites remained for analysis, as summarized in Table 1. This filtered dataset allows us to study how the representation evolves as the distance threshold *t* varies, while ensuring that both the structural and compositional descriptors are consistently defined across the analyzed binding sites.

### 2.3 TOUGH-C1 dataset

The TOUGH-C1 dataset [42] provides a larger benchmark derived from protein–ligand complexes in the PDB. It contains binding sites associated with nucleotide-like ligands and heme, together with a control set of binding sites not associated with these ligand classes. The control set was constructed to ensure low sequence and structural similarity to the other groups, resulting in a chemically and structurally distinct collection of binding sites. A summary of the original and filtered datasets, including the number of binding sites in each group, is provided in Table 2.

**Table 2:** Composition of the TOUGH-C1 dataset used in this study.

| Binding site type | Original dataset | Analyzed dataset |
| --- | --- | --- |
| Nucleotide-binding sites | 1,553 | 1,547 |
| Heme-binding sites | 596 | 592 |
| Control sites | 1,946 | 1,863 |

We use TOUGH-C1 to evaluate FUSED in two binary classification settings: nucleotide-binding sites versus control sites and heme-binding sites versus control sites. These tasks provide a larger-scale complement to the multiclass EK analysis and allow us to assess how well the representation distinguishes specific binding-site classes from a broader background.

### 2.4 TOUGH-M1 dataset

The TOUGH-M1 benchmark, introduced by [16], consists of approximately one million pairs of protein–ligand binding sites obtained from the PDB. The dataset was constructed specifically from complexes involving drug-like ligands: only complexes whose bound ligand has a Tanimoto coefficient of at least 0.5 to an FDA-approved drug were retained. To reduce redundancy, protein complexes were first clustered by sequence similarity, after which representative structures binding chemically diverse ligands were selected from each cluster. The bound ligands were then clustered according to chemical similarity. Positive pocket pairs consist of globally dissimilar proteins that bind chemically similar ligands from the same ligand cluster, whereas negative pairs consist of globally dissimilar proteins whose ligands belong to different clusters. This procedure yielded 505,116 positive and 556,810 negative pocket pairs in the final TOUGH-M1 dataset.

We use TOUGH-M1 to evaluate whether FUSED can support pocket matching for pharmacologically relevant, drug-like binding sites. In contrast to the multiclass classification task considered for EK and the two binary classification tasks considered for TOUGH-C1, TOUGH-M1 poses a pairwise matching problem. Given two binding sites, the objective is to determine whether they are likely to bind chemically similar ligands.

Of the 7,524 pockets in the official TOUGH-M1 pocket list, 7,477 had usable FUSED feature vectors. Of these, 7,447 also had Research Collaboratory for Structural Bioinformatics Protein Data Bank (RCSB PDB) 30% sequence-identity cluster assignments and were retained for the cluster-based neural-network evaluation. Pairs involving excluded pockets were removed from the analysis, resulting in the pocket and pair counts shown in Table 3.

**Table 3:** TOUGH-M1 pocket and pair counts used in the study.

| Quantity | Official TOUGH-M1 | Analyzed in this study |
| --- | --- | --- |
| Pockets | 7,524 | 7,447 |
| Positive pairs | 505,116 | 498,628 |
| Negative pairs | 556,810 | 540,764 |
| Total pairs | 1,061,926 | 1,039,392 |

## 3 Structural and Compositional Descriptors

In this section, we construct structural and compositional descriptors of ligand binding sites defined at a given distance threshold. As the threshold *t* varies, the set of atoms defining the binding site changes, and our goal is to capture how the structural and chemical characteristics of the binding site depend on this choice of *t*.

A central requirement for these descriptors is that they be defined individually for each binding site, without reliance on pairwise comparisons or alignment-based measures of similarity. The more difficult part of this construction is obtaining a structural descriptor that is comparable across binding sites despite arbitrary orientations and the absence of a common coordinate system. To address this, we represent structure using the Covariance of Distances to Principal Axes (CDPA) [41], which provides a coordinate-invariant description of binding site geometry. However, CDPA contains no compositional information, so we complement it with descriptors based on the elemental composition of the atoms in the binding site. Together, these provide a representation that reflects both the spatial structure and chemical composition of the binding site.

### 3.1 Structural Descriptors via CDPA

CDPA addresses the arbitrary coordinate system in PDB files by encoding structural information relative to the three principal axes of variation of the binding site. For a fixed distance threshold *t*, let **P**_*i*_(*t*) denote the *i*^th^ binding site at threshold *t*, defined as in Section 2.1 as the set of protein atoms within distance *t* of the ligand. Principal component analysis is applied to the atomic coordinates in **P**_*i*_(*t*), yielding three orthogonal principal axes that define a coordinate system determined by the geometry of the binding site.

We measure the distances between each atom and the three principal axes. For an atom, the distance to an axis is its Euclidean distance to its projection onto that axis. Let *d*_*aj,t*_ denote the distance of the *a*^th^ atom to the *j*^th^ principal axis, where *a* = 1, …, *n*_*i,t*_, *j* = 1, 2, 3, and *n*_*i,t*_ is the number of atoms in **P**_*i*_(*t*). Let

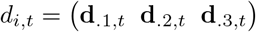

denote the *n*_*i,t*_ × 3 matrix of distances, where **d**_.*j,t*_ is the vector of distances to the *j*^th^ principal axis.

The covariance matrix

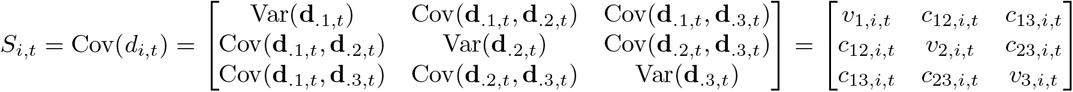

summarizes how the atoms in **P**_*i*_(*t*) are distributed relative to the principal axes. The diagonal elements *v*_1,*i,t*_, *v*_2,*i,t*_, *v*_3,*i,t*_ measure the spread of atoms around each axis, while the off-diagonal elements capture the joint variation between these distances. Together, these quantities describe the overall shape and spatial organization of the binding site at threshold *t* in a manner that is invariant to rotation and translation.

We vectorize *S*_*i,t*_ as

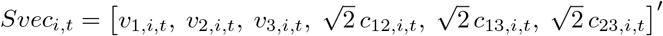

to obtain a six-dimensional feature vector that serves as the structural descriptor of the binding site at threshold *t*. This allows us to work with standard multivariate representations rather than treating *S*_*i,t*_ as a matrix-valued object, thereby avoiding the need to define distances or metrics between covariance matrices. The resulting vector *Svec*_*i,t*_ can be directly used in subsequent analyses and combined with compositional descriptors. The off-diagonal terms are scaled by 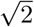 so that *Svec*_*i,t*_ preserves the Frobenius norm of *S*_*i,t*_, ensuring that the relative contributions of the variance and covariance terms are retained in the vectorized representation.

### 3.2 Compositional Descriptors via ILR Transformation

While CDPA captures the geometric structure of the binding site, it does not account for elemental composition, which provides complementary information about the atoms involved. To incorporate this information, we summarize the relative abundances of selected atom types within **P**_*i*_(*t*), the *i*^th^ binding site at threshold *t*.

More generally, suppose that *m* atom types are considered and let

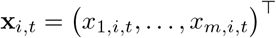

denote the corresponding compositional vector. Because these compositions lie in the simplex of nonnegative vectors whose components sum to 1, standard Euclidean methods are not directly applicable, as they do not respect this constraint. We therefore transform them into Euclidean space using the isometric log-ratio (ILR) transformation [9], which produces orthonormal coordinates and preserves the Aitchison geometry of the simplex.

The ILR transformation of **x**_*i,t*_ yields *m* − 1 coordinates given by

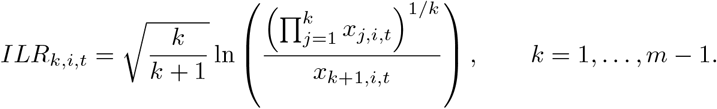

For the analyses considered in this paper, we use three atom types: carbon (C), oxygen (O), and nitrogen (N). Accordingly, let

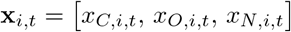

denote their proportions within **P**_*i*_(*t*). The resulting two ILR coordinates are

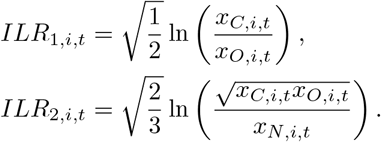

The vector [*ILR*_1,*i,t*_, *ILR*_2,*i,t*_]therefore provides a two-dimensional Euclidean representation of elemental composition at threshold *t*.

The choice of C, O, and N reflects the elemental distributions in the datasets analyzed here. Non-hydrogen atoms other than C, O, and N represented only a small fraction of each binding site, with median site-level proportions of 0.41% for Extended Kahraman and 0.48% for TOUGH-C1. Such sparse components frequently produce zero or very small proportions, which are problematic for ILR coordinates because the logarithms are undefined at zero and can become unstable for very small values. We therefore use C, O, and N as the compositional basis for the present analyses. The general ILR formulation above nevertheless allows additional atom types to be incorporated in applications where they are sufficiently represented.

### 3.3 Combined Structural and Compositional Representation

The structural and compositional descriptors are combined to form a unified representation of the binding site at threshold *t*. Specifically, for the *i*^th^ binding site and threshold *t*, we define the eight-dimensional feature vector

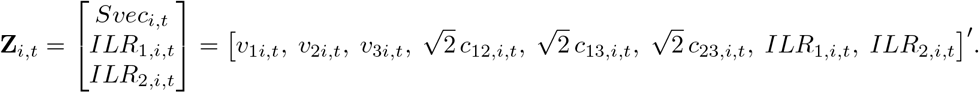

This representation integrates geometric and elemental information into a single Euclidean feature vector. The first six components capture the spatial distribution of atoms relative to the principal axes via CDPA, while the final two components encode the elemental composition through the ILR transformation. Together, these features provide a more comprehensive description of the binding site at threshold *t* than either alone, enabling direct comparison across binding sites using standard multivariate techniques.

## 4. Evaluating Representation Adequacy

In this section, we introduce simple diagnostic tools used to evaluate whether a representation captures distinguishing information among ligand groups.

### 4.1 Evaluation Framework

We now examine how the descriptors introduced in Section 3 characterize within-group variability and between-group differences among ligand binding sites. In particular, we assess whether the CDPA and ILR representations, both individually and jointly, capture structure that makes patterns of variability within each group apparent while also distinguishing among ligand classes. To evaluate these aspects, we use both graphical tools and simple classification studies as complementary diagnostics. The graphical representations provide direct insight into the distribution and spread of the data, while classification results summarize how well these patterns translate into distinguishability between groups. We carry out this assessment at multiple distance thresholds to examine how both within-group variability and between-group separation depend on the choice of threshold used to define the binding sites.

To directly examine the within-group variability and between-group differences described above, we begin with graphical diagnostics by visualizing the representations across ligand groups. For the compositional features (ILR coordinates), we examine scatterplots of the full feature space, while for the structural and combined representations, we consider principal component projections to capture the dominant modes of variation. In each case, binding sites are colored according to ligand group, allowing us to assess separation between groups as well as the spread and heterogeneity within groups. These visualizations provide direct insight into how the representation organizes the data, highlighting patterns of overlap, dispersion, and clustering that are not fully captured by scalar performance measures.

To complement these visual assessments, we use classification as a diagnostic tool to evaluate whether the observed structure translates into consistent distinguishability between ligand groups. More specifically, we consider three statistical learning methods that perform classification using fundamentally different frameworks, allowing us to assess the extent to which performance depends on the classification procedure versus the underlying structure of the representations.

The first method is the nearest Mahalanobis mean classifier, which assigns each LBS to the group whose mean is closest under a covariance-adjusted distance. This can be viewed as a nearest-centroid rule that accounts for differences in scale and correlation among features, providing a geometry-based classification approach that reflects both location and variability within each group. The second method is multinomial logistic regression, which models the probability that each LBS belongs to each ligand class using a linear function of the features and assigns observations based on the largest predicted probability. This method assesses whether the representation supports linear separation among groups. The third method is random forest, an ensemble of decision trees that produces classifications through recursive partitioning and aggregation. In contrast to the other two methods, it is able to capture nonlinear relationships and interactions among features. Details for each of these methods can be found in the Appendix.

We used repeated stratified 5-fold cross-validation with *k* repetitions to evaluate classification performance, where *k* denotes the number of repetitions used in the corresponding analysis. In each repetition, the data were randomly divided into five folds while preserving the ligand group proportions as closely as possible. Each fold was then used once as the held-out test fold, with the remaining four folds used for model fitting. This produced *k* × 5 held-out evaluations. Reported accuracies are the mean and standard deviation of the *k* × 5 held-out fold accuracies. For comparisons among representations or modeling conditions, we used the paired sign-flip randomization test for the fold-wise accuracy differences. This paired sign-flip procedure is a form of Fisher permutation test [23]. The pairing was defined by the same repetition and fold. Each representation or modeling condition was evaluated on the same held-out fold. The performance measure from one condition was then paired with the corresponding performance measure from the other condition on that identical fold. Each *p*-value tests whether the average difference in accuracy between the two representations or modeling conditions being compared is different from zero.

### 4.2 Dependence on Distance Thresholds

We examine how the CDPA and ILR representations vary with the choice of distance threshold, focusing on changes in within-group variability and between-group separation at two representative values, 5.3 Å and 12.0 Å. The CDPA-based structural descriptors shown in Figure 1 exhibit clear separation among several ligand groups at 5.3 Å, consistent with previously reported results for this dataset. At 12.0 Å, the representation changes in a non-uniform way: while separation between some groups decreases, others, most notably FAD, become more clearly separated, and within-group variability increases for certain ligands such as PO_4_. These structural patterns are reflected in the classification performance displayed in Table 4, where CDPA alone provides substantially higher classification accuracy than ILR alone at both thresholds, although performance declines at 12.0 Å in line with the increased overlap observed for many groups. Together, these results indicate that CDPA captures meaningful structural information across thresholds, but that the nature of this information changes with *t*.

**Table 4:**
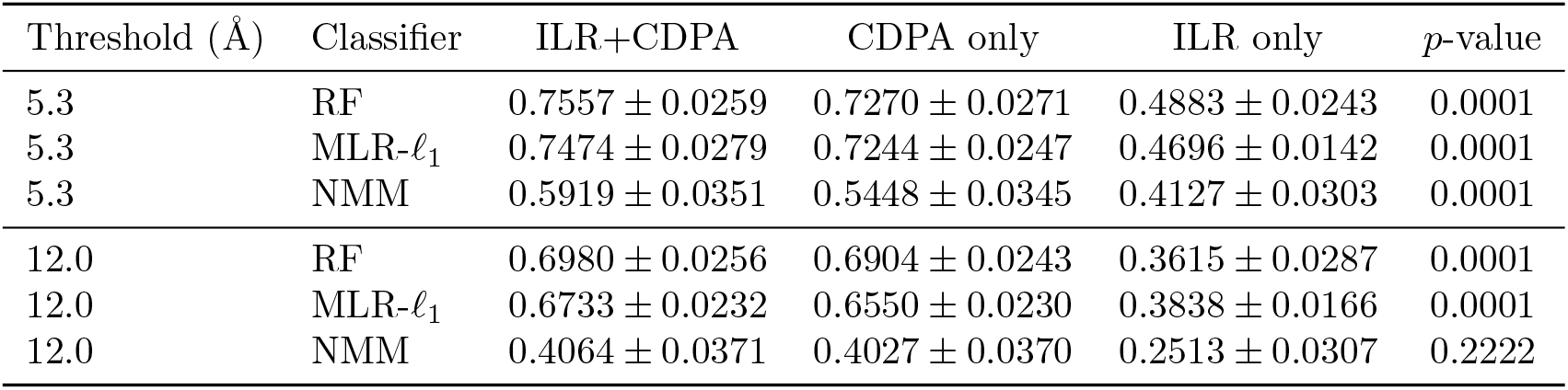
Classification performances for different representations at distance thresholds 5.3 Å and 12.0 Å for the Extended Kahraman dataset.Accuracies are reported as mean ± standard deviation over 100 fold evaluations from 20 repeated 5-fold cross-validation. The *p*-value is the result of performing a paired sign-flip randomization test comparing the difference between ILR+CDPA and CDPA-only accuracies. Here, RF denotes random forest, MLR-*l*_1_ denotes multinomial logistic regression with *l*_1_ penalty, and NMM denotes the nearest Mahalanobis mean classifier.

| Threshold (Å) | Classifier | ILR+CDPA | CDPA only | ILR only | $p$-value |
| --- | --- | --- | --- | --- | --- |
| 5.3 | RF | $0.7557 \pm 0.0259$ | $0.7270 \pm 0.0271$ | $0.4883 \pm 0.0243$ | 0.0001 |
| 5.3 | MLR-$\ell_1$ | $0.7474 \pm 0.0279$ | $0.7244 \pm 0.0247$ | $0.4696 \pm 0.0142$ | 0.0001 |
| 5.3 | NMM | $0.5919 \pm 0.0351$ | $0.5448 \pm 0.0345$ | $0.4127 \pm 0.0303$ | 0.0001 |
| 12.0 | RF | $0.6980 \pm 0.0256$ | $0.6904 \pm 0.0243$ | $0.3615 \pm 0.0287$ | 0.0001 |
| 12.0 | MLR-$\ell_1$ | $0.6733 \pm 0.0232$ | $0.6550 \pm 0.0230$ | $0.3838 \pm 0.0166$ | 0.0001 |
| 12.0 | NMM | $0.4064 \pm 0.0371$ | $0.4027 \pm 0.0370$ | $0.2513 \pm 0.0307$ | 0.2222 |

**Figure 1:**
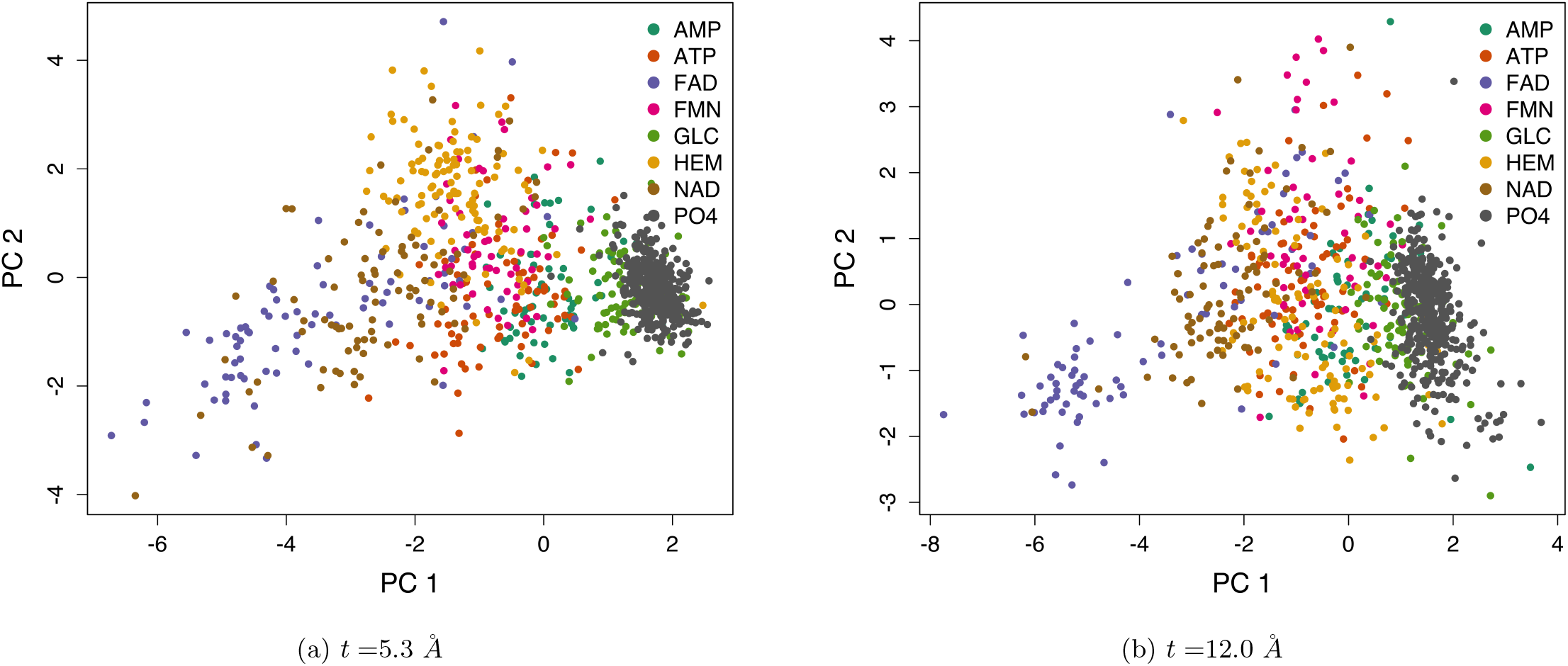
Two-dimensional PCA score plots based on the six CDPA components at two distance thresholds. Each point represents an LBS and is colored by ligand group.

In contrast, the ILR-based compositional descriptors displayed in Figure 2 show substantially weaker separation at both thresholds, with considerable overlap among ligand groups even at 5.3 Å. This limited separation is directly reflected in classification performance, where ILR alone performs substantially worse than CDPA, though still above chance, indicating the presence of some compositional signal. At 12.0 Å, the ILR representations become more tightly clustered, and classification performance deteriorates further, consistent with the reduced distinction between groups. These results show that while compositional information captures certain aspects of the binding environment, it provides substantially weaker discrimination than CDPA alone in the EK dataset, with performance also varying with the choice of threshold.

**Figure 2:**
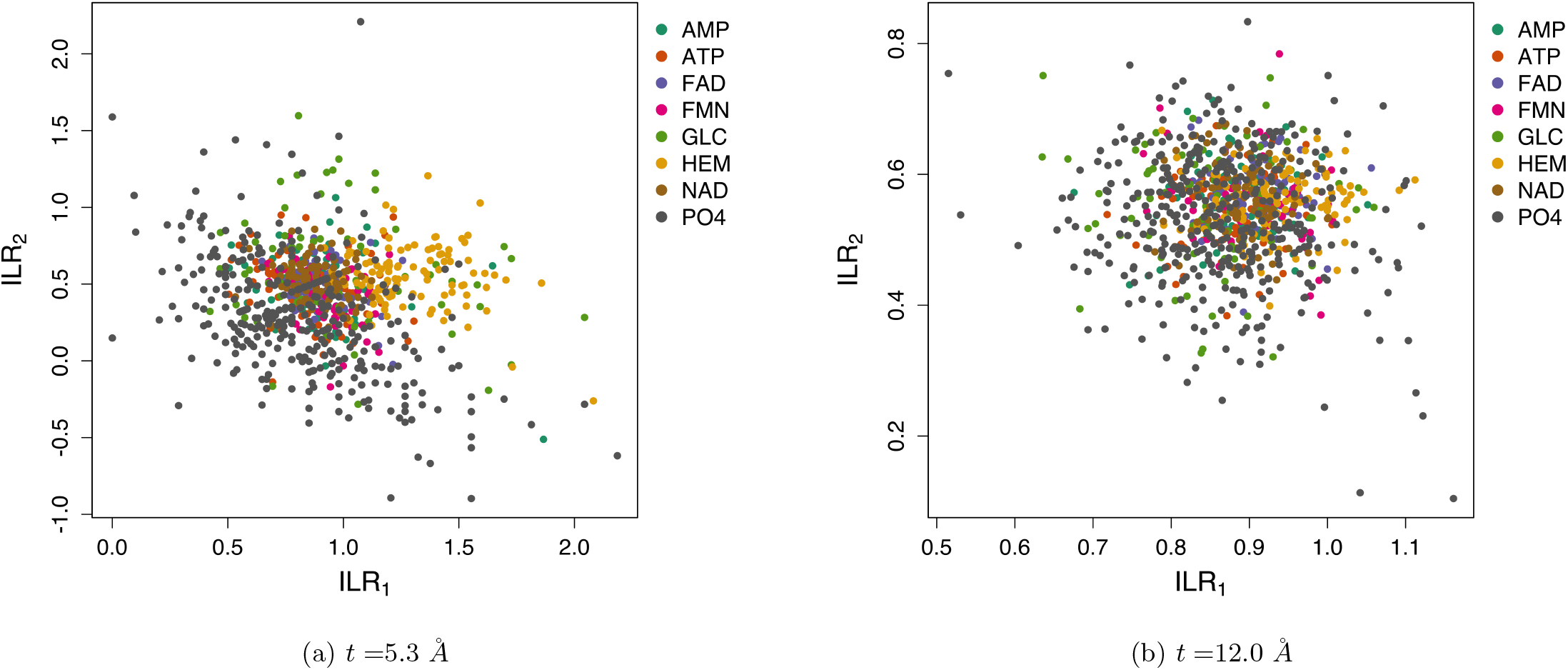
Two-dimensional scatter plots of the ILR-transformed compositional features at two distance thresholds. Each point represents an LBS and is colored according to ligand group.

When CDPA and ILR features are combined, as in Figure 3, the resulting representation exhibits clearer separation for certain ligand groups, particularly HEM, at 5.3 Å, indicating that compositional information provides additional discriminatory structure beyond CDPA alone. This improvement is reflected in classification performance, where all methods show increased accuracy relative to CDPA alone at this threshold. At 12.0 Å, however, the combined representation closely resembles CDPA, and the corresponding classification gains are generally smaller, consistent with the diminished contribution of compositional information at larger thresholds. Notably, random forest and multinomial logistic regression achieve broadly similar performance across settings, suggesting that the results are due largely to the observed discriminatory structure rather than the specific linear or nonlinear classification procedure. The nearest Mahalanobis mean classifier shows a larger relative gain from the addition of ILR at 5.3 Å, suggesting that the added compositional information alters the geometry of the feature space in a way that particularly benefits distance-based classification.

**Figure 3:**
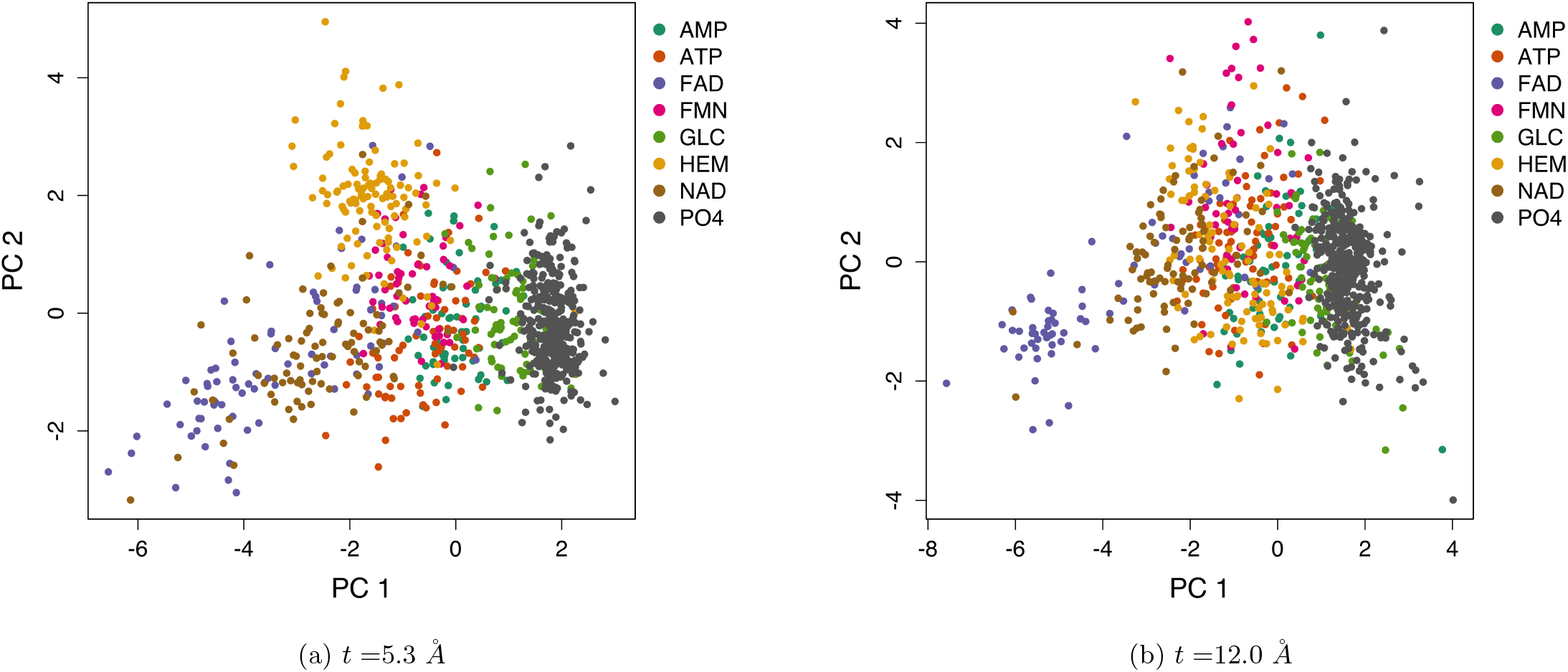
Two-dimensional PCA score plots based on the combined ILR and CDPA features at two distance thresholds.

**Figure 4:**
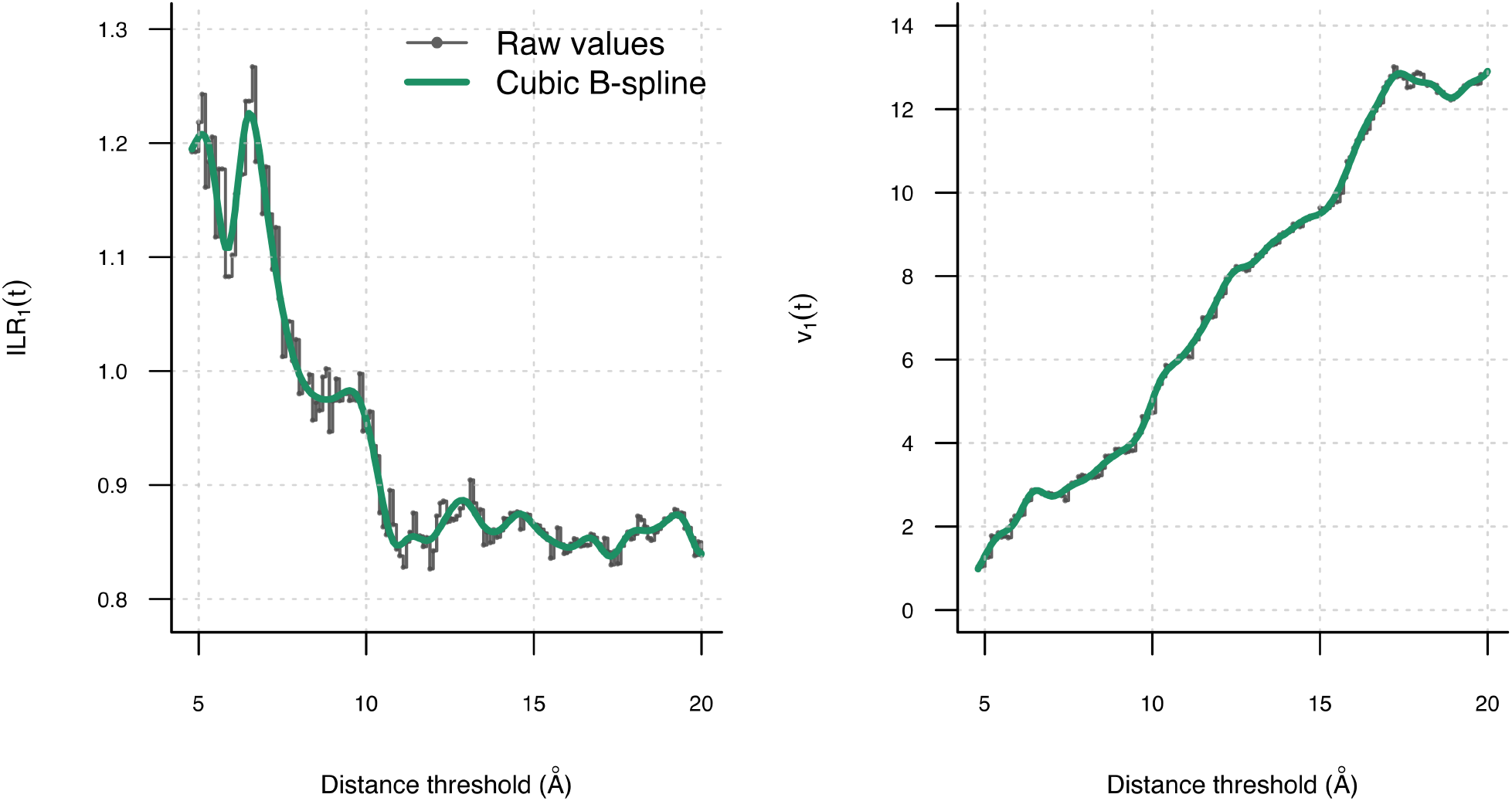
Illustration of *ILR*_1_(*t*) and *v*_1_(*t*) and their fitted cubic B-spline representations for one representative HEM binding site from the EK dataset (PDB ID: 2G5G).

Taken together, the visual and classification results provide a consistent picture of how the representations behave across thresholds. While classification performance summarizes overall discriminative ability in a way that visualization alone cannot, it does not fully capture the degree or geometry of separation between ligand groups observed in the visualizations. For example, certain ligand groups such as FAD exhibit increased separation at larger thresholds without corresponding improvements in classification accuracy. This indicates that additional structural information may be present in the representation but is not fully reflected in aggregate performance measures. Overall, these results demonstrate that structural and compositional features provide complementary information, but neither representation alone yields consistently strong separation among ligand groups at a single threshold. Different choices of *t* reveal different aspects of the binding site structure, and neither visual separation nor classification performance alone fully characterizes these changes. This motivates the development of representations that incorporate information across multiple distance thresholds.

## 5 Functional Unification of Structural and Elemental Descriptors (FUSED)

The Functional Unification of Structural and Elemental Descriptors (FUSED) representation models each ligand binding site as a multivariate function over the distance threshold, combining structural dispersion and chemical composition.

### 5.1 FUSED Representation

We now formalize the functional representation underlying FUSED. Let *T* ⊂ ℝ_+_ denote the domain of distance thresholds. For each ligand binding site *i* = 1, …, *N*, recall that at each threshold *t* ∈ *T* we obtain an eight-dimensional feature vector **Z**_*i,t*_ combining structural and compositional information. We define the functional representation of the *i*^th^ binding site by

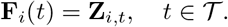

Thus,

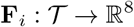

is a multivariate functional object whose components correspond to the structural variance and covariance summaries from CDPA together with the ILR-transformed compositional features.

The threshold domain *T* may be chosen in different ways depending on the scientific question. It may consist of a single threshold, a finite set of thresholds, or an interval of thresholds. A single threshold gives a cross-sectional representation of the binding site at one neighborhood size, whereas an interval allows the representation to describe how structural and compositional features evolve as progressively more distant atoms are included.

### 5.2 Implementation of FUSED over a Distance Interval

When *T* = [*t*_min_, *t*_max_] is an interval, the binding site **P**_*i*_(*t*) and the resulting descriptor vector **Z**_*i,t*_ are evaluated in practice on a dense grid of threshold values rather than at every *t* ∈ T. In addition, because membership in **P**_*i*_(*t*) changes only when the threshold crosses the distance of an additional protein atom from the ligand, the resulting threshold-dependent descriptors are inherently stepwise functions of *t*. We therefore use smooth basis expansions to obtain functional approximations to these grid-evaluated, stepwise descriptor trajectories. The resulting smooth functions should be viewed as representations of the threshold-dependent descriptors rather than as an assumption that the underlying atom-inclusion process itself varies continuously.

For the analyses in this paper, we evaluate the descriptors on a grid with spacing 0.1 Å over the interval [*t*_min_, *t*_max_]. Each component of **F**_*i*_(*t*) is then represented using a cubic B-spline basis expansion with penalized least squares. Specifically, for each component *k* = 1, …, 8,

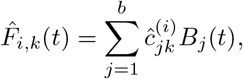

where *B*_*j*_(*t*) are B-spline basis functions, 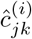 are the estimated coefficients for the *k*^th^ component of the *i*^th^ binding site, and *b* is the number of basis functions. A common smoothing parameter *λ* = 10^*−*3^ is used across all components. The number of basis functions *b* is selected from the candidate set {*M/*2, *M/*3, *M/*4, *M/*5}, where *M* is the number of grid points in [*t*_min_, *t*_max_]. We choose the smallest admissible value of *b* with *b* ≥ 4 to ensure a stable functional representation as *M* varies with the length of the interval [49]. The value *λ* = 10^*−*3^ was used as the reference smoothing parameter in the main analyses. To assess sensitivity to this choice, we also considered *λ* = 10^*−*4^, 10^*−*2^, and 10^*−*1^. Downstream classification performance was highly stable across these values (Table A1), indicating that the analyses are not materially sensitive to the degree of smoothing over the range considered.

### 5.3 Uses of the FUSED Representation

FUSED provides a representation of binding-site structure and composition across the threshold domain rather than prescribing a particular downstream analysis. Consequently, the resulting functions can be used in different ways depending on the scientific or statistical objective. They may be examined directly to visualize and explore threshold-dependent patterns, summarized through functional dimension-reduction methods such as MFPCA, or used as input for subsequent statistical and machine-learning procedures. The representation can also be restricted to particular thresholds or ranges of thresholds when the objective is to study where relevant differences occur.

The remainder of the paper illustrates these complementary uses of FUSED. We first examine the functional representations directly and then consider MFPCA as one approach for obtaining lower-dimensional summaries. We subsequently investigate the role of the threshold domain and evaluate the representation in downstream classification and pocket-matching tasks. These analyses are intended to demonstrate different ways in which FUSED can be used rather than to define additional components of the representation itself.

## 6 Exploratory and Descriptive Analysis Using FUSED

The functional nature of FUSED provides several ways to examine how binding-site characteristics vary across the distance domain. We first examine the individual functional components directly to characterize between-group and within-group patterns as the distance threshold changes. We then use multivariate functional principal component analysis (MFPCA) to obtain low-dimensional summaries of the joint variation across the structural and compositional components. Together, these analyses illustrate complementary ways of exploring the information contained in the FUSED representation before considering its use in subsequent statistical and machine-learning procedures.

### 6.1 Direct Analysis via the Functional Representation

We use the functional representation to examine how structural and compositional variation in ligand binding sites evolves across distance thresholds. As in Section 2, we focus on the extended Kahraman (EK) dataset, which provides a consistent and well-characterized collection of ligand binding sites for studying these patterns. We consider a broad domain of distance thresholds *T* = [*t*_min_, *t*_max_], deliberately chosen to avoid excluding potentially informative regions. This allows us to assess how information in the data changes as the neighborhood around the ligand expands, rather than restricting attention to a fixed threshold a priori.

We first examine between-group variation through the group mean functions for each component of the representation. These reveal clear group-level differences, particularly at smaller distance thresholds. For the compositional components, such as ILR_1_(*t*) and ILR_2_(*t*), the group mean functions exhibit substantial separation near the ligand, indicating that chemical composition differs meaningfully across ligand groups at short distances. However, as the distance threshold increases, these mean functions tend to converge, suggesting that compositional differences become less pronounced farther from the ligand, as shown in Figure 5.

**Figure 5:**
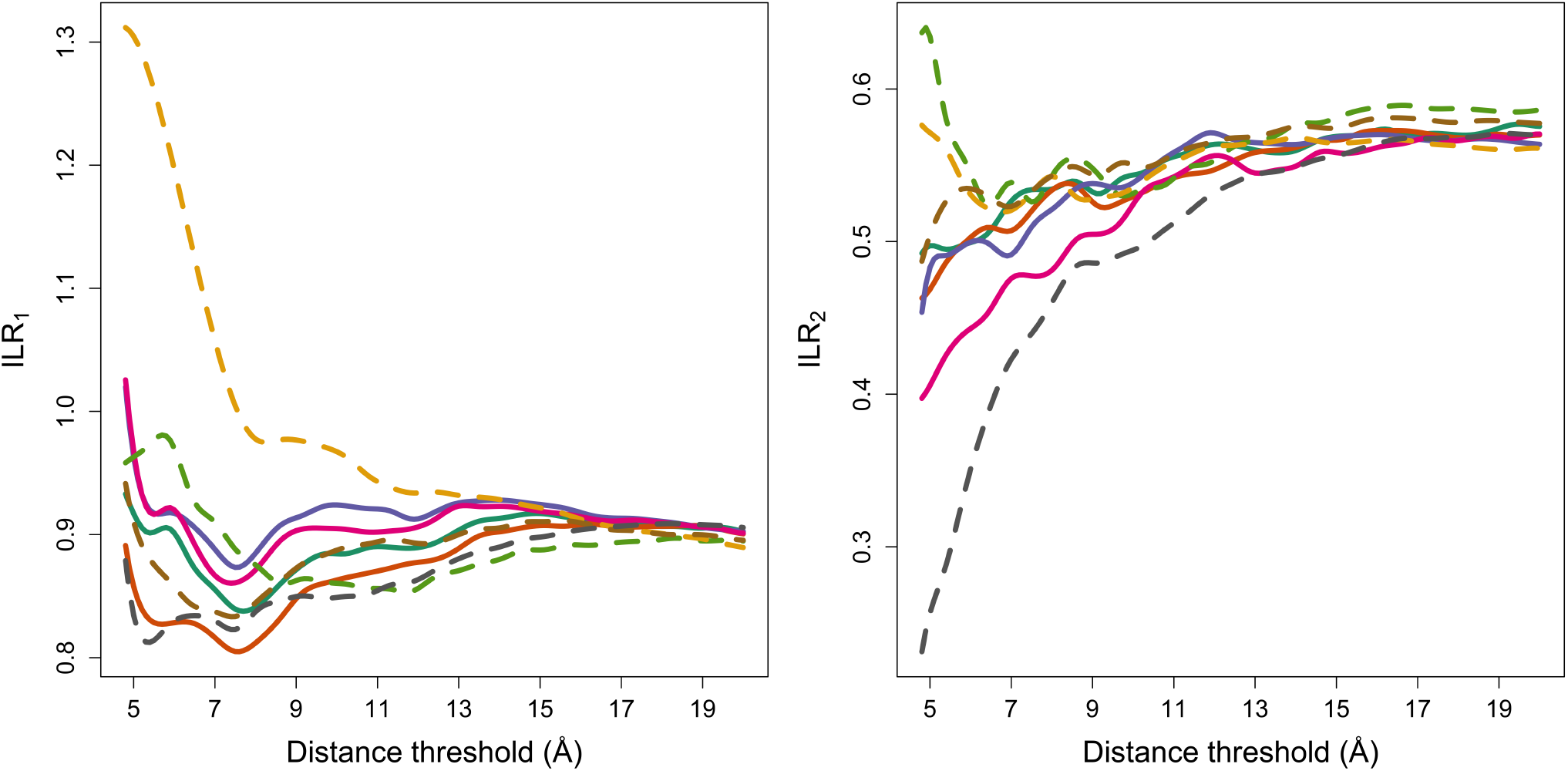
Group mean functions of ILR_1_ and ILR_2_ as a function of distance threshold. Line colors and styles indicate ligand groups: AMP, ATP, FAD, FMN, GLC, HEM, NAD, PO_4_.

In contrast, the structural components derived from CDPA exhibit a different pattern. The variance functions *v*_*k*_(*t*) and several covariance functions *c*_*ij*_(*t*) maintain more persistent separation across ligand groups over a larger portion of the domain. At the same time, certain covariance components display nontrivial changes in their relative behavior as the distance threshold increases, indicating that the joint variation in distances to the principal axes evolves with the inclusion of more distant atoms, as in Figures 6 and 7. These observations highlight that different components of the representation capture distinct aspects of variability across the domain *T*.

**Figure 6:**
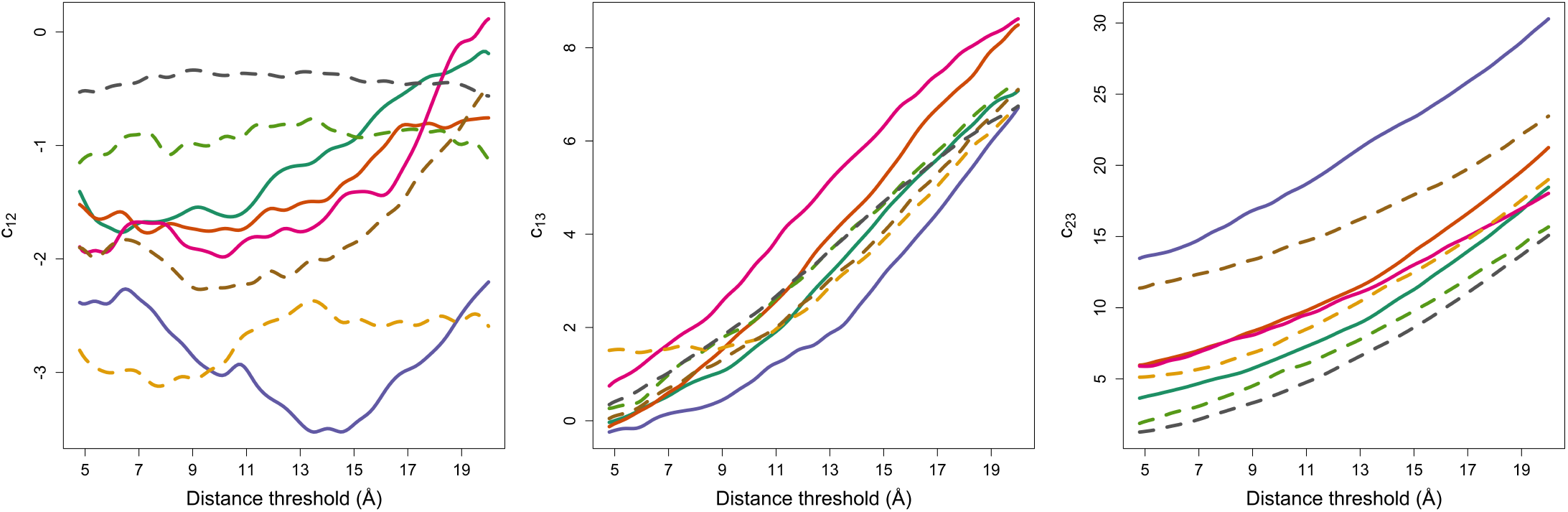
Group mean functions of *c*_12_, *c*_13_, and *c*_23_ as a function of distance threshold. Line colors and styles indicate ligand groups: AMP, ATP, FAD, FMN, GLC, HEM, NAD, PO_4_.

**Figure 7:**
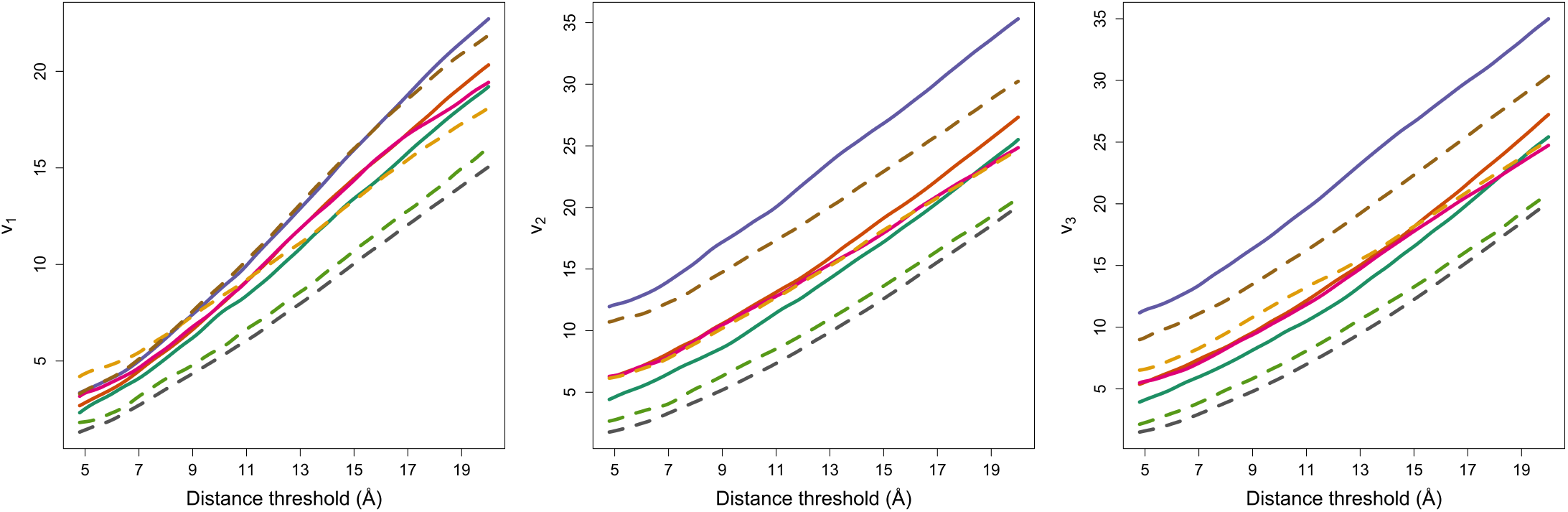
Group mean functions of *v*_1_, *v*_2_, and *v*_3_ as a function of distance threshold. Line colors and styles indicate ligand groups: AMP, ATP, FAD, FMN, GLC, HEM, NAD, PO_4_.

To better understand how these between-group patterns relate to variability within ligand groups, we next examine the functional behavior of individual binding sites within a representative group in Figure 8. For the compositional components, we observe substantial within-group variability at small distance thresholds, particularly near the ligand. This variability decreases markedly as the distance threshold increases, with curves becoming more stable and similar across sites. When viewed alongside the group mean behavior, this indicates that although strong between-group differences exist at small distances, they are accompanied by substantial within-group variation. This helps explain why such regions may not necessarily yield consistent discrimination when used directly in downstream tasks.

**Figure 8:**
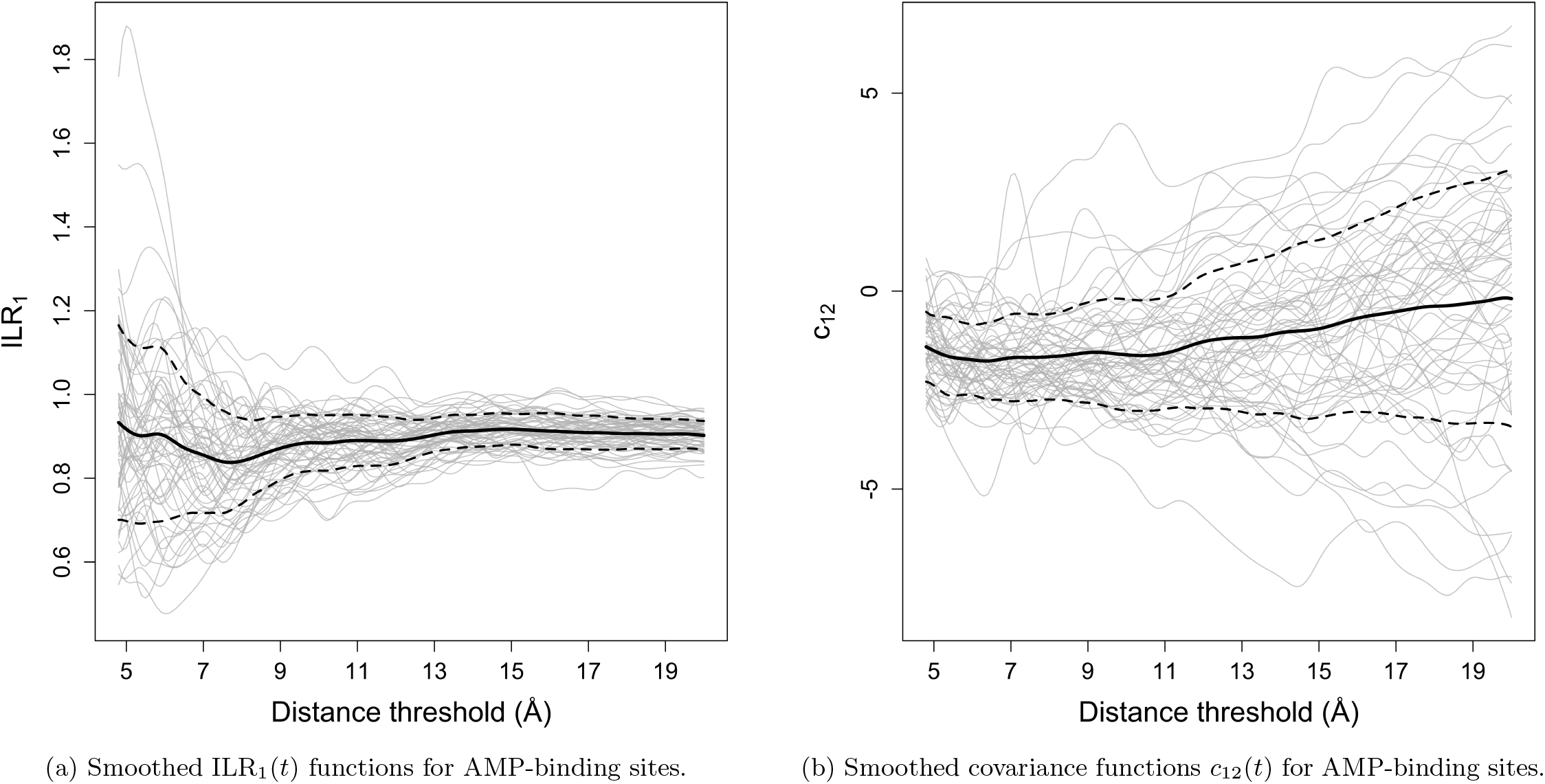
Functional representations of AMP-binding sites with ILR_1_(*t*) and *c*_12_(*t*) functions. Grey curves represent individual binding sites, the solid black curve is the mean function, and dashed curves indicate mean ± standard deviation.

At larger distance thresholds, both the between-group differences and the within-group variability for the compositional components diminish, so the group structure becomes less apparent from these visual summaries alone. This does not imply that the corresponding regions contain no useful discriminatory information, since differences that are difficult to distinguish visually may still be detected or exploited through multivariate statistical and machine-learning procedures. Rather, these plots show that the compositional differences are most visually pronounced near the ligand and become increasingly subtle as the threshold increases. The analyses that follow therefore complement direct visualization with dimension reduction and classification-based assessments of how much useful information remains across the threshold domain.

For the structural components, the relationship between within-group variability and between-group differences is more stable across the domain. The variance functions and several covariance functions exhibit clearer and more persistent differences between groups, with comparatively structured within-group behavior. However, not all structural components behave uniformly: some covariance functions show evolving patterns over distance, suggesting that additional information may be gained by considering how these functions change rather than relying on a single fixed threshold.

Taken together, these observations indicate that the functional representation provides a rich but highly complex description of ligand binding sites. It simultaneously captures variation across ligand groups, variability within groups, and evolving relationships among multiple components as the distance threshold varies. While the preceding plots allow us to examine these aspects individually, it is difficult to synthesize them into a coherent global picture or to assess their combined effect when considered at this level of detail.

This motivates the need for a principled dimension reduction approach that can jointly account for variation across all components and across the domain *T*. Such an approach allows us to move from a detailed but unwieldy representation to a lower-dimensional summary that preserves the key sources of variability while providing a more manageable summary of the overall structure of the data for subsequent analysis.

### 6.2 Visualization in Low Dimensions via MFPCA

The functional representation introduced above provides a detailed description of ligand binding sites, capturing variation across multiple features and across the full domain *T*. Because this information is distributed across several functional components and distance thresholds, dimension reduction can be useful for obtaining lower-dimensional summaries of the dominant patterns of variation. A variety of dimension-reduction approaches could be applied to the FUSED representation depending on the objective of the analysis. For exploratory visualization of multivariate functional data, we use multivariate functional principal component analysis (MFPCA), which extends both multivariate principal component analysis and univariate functional principal component analysis to multivariate functional data [6]. This allows us to summarize and visualize the joint variation of the ILR and CDPA functions rather than examining each component or threshold separately.

Technical details of the MFPCA procedure, including the functional formulation, covariance structure, and eigenfunction decomposition, are provided in Appendix B. A practical consideration specific to the FUSED representation is the relative scaling of its different functional components. As is apparent in Figures 5– 7, the ILR and CDPA functions can differ substantially in numerical scale. Without adjustment, components with substantially larger variances can dominate the covariance structure used by MFPCA, causing the leading principal components to reflect primarily differences in marginal scale rather than the joint patterns of variation across the functional components. We therefore rescale the ILR channels by constants so that their variability is on a scale more comparable to that of the CDPA functions, allowing the joint covariance structure across components to potentially play a greater role in the resulting MFPCA. The specific scaling procedure used here is also described in Appendix B.

MFPCA is then applied to the collection of functions for each binding site, producing principal component scores that summarize the dominant patterns of joint variation across the components of FUSED and across *T*. These scores provide low-dimensional views of variation that would be difficult to assess from examining the eight functional components simultaneously. By plotting the leading MFPC scores and identifying points by ligand group, we can examine whether the dominant sources of variation in FUSED also reveal group-level structure, including patterns of separation, overlap, and within-group heterogeneity.

Figure 9 illustrates how MFPCA can be used to visualize the dominant patterns of joint variation in the FUSED representation for the EK dataset. Over the shorter interval 4.8–6.0 Å, several group-level patterns are already apparent. PO_4_ is clearly separated from most of the other ligand groups, although some overlap with GLC is present. FAD and NAD are also substantially separated from the remaining groups, while showing some overlap with one another. In contrast, the remaining ligand groups occupy largely overlapping regions of the first two MFPC score dimensions.

**Figure 9:**
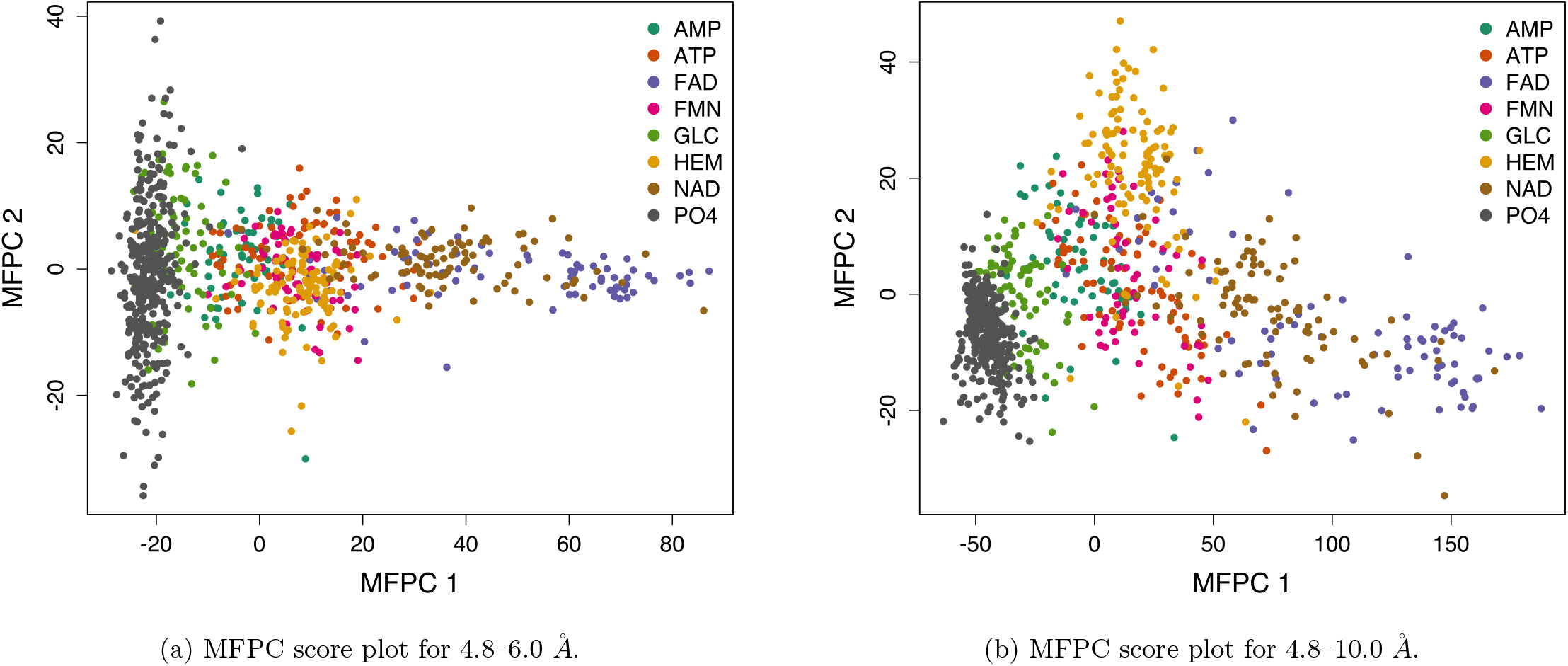
Two-dimensional scatter plots of the first two MFPC scores of the joint functional representation based on the ILR and CDPA channels of the EK dataset, shown for two different upper distance-threshold limits.

When the domain is extended to 4.8–10.0 Å, the same broad patterns remain, but additional distinctions become visible. PO_4_ and GLC show clearer separation, as do NAD and FAD, although some overlap remains in the latter case. HEM also becomes more distinct from the groups that remain substantially overlapping in this two-dimensional projection. Thus, for the EK dataset, extending the threshold domain incorporates additional functional information that reveals group-level structure not as clearly visible when the representation is restricted to the shorter interval. This illustrates an important exploratory advantage of the functional representation: examining different portions of the distance domain can reveal group-level distinctions in the leading MFPC scores that are less apparent when the representation is restricted to a narrower range of thresholds.

## 7 Impact of the Distance Interval on Analysis

The exploratory analyses in Section 6 demonstrate that the information revealed by FUSED can change as different portions of the distance interval are included in the representation. This raises an important practical consideration for the use of FUSED: the choice of distance interval can affect the information available for subsequent analysis. Moreover, there is no reason to expect a single interval to be equally informative for all purposes. The useful range of thresholds may depend on the dataset, the objective of the analysis, and the particular analytical procedure being applied. We illustrate these effects here, with particular emphasis on classification as a setting in which they can be quantified directly. The classification studies in the following sections then address interval selection separately for the corresponding datasets, tasks, and procedures.

We first consider the EK dataset, where the direct functional analyses provide insight into how different sources of information change as the interval is extended. The ILR group mean functions in Figure 5 show the strongest compositional differences at shorter distances, followed by increasing convergence among the groups as the threshold increases. In contrast, the CDPA variance functions in Figure 7 maintain group differences over a broader range of thresholds. The covariance functions in Figure 6 exhibit still different behavior, with some group relationships changing as increasingly distant atoms are included. Thus, extending the distance interval does not simply add more of the same information: the relative contributions and behavior of different components of FUSED change across the interval.

The effect of these changes can also be explored through classification. Figure 10 shows classification accuracy for the EK dataset as the lower endpoint is held fixed at 4.8 Å and the upper endpoint *t*_max_ is varied. These results are based on a simpler cross-validation scheme used for exploratory illustration rather than the nested cross-validation procedure used for the subsequent classification analyses. Accordingly, the individual accuracies in this figure are not used as final estimates of predictive performance or to select the intervals used later in the paper. Instead, the figure illustrates how the apparent discriminatory utility of the FUSED representation changes as increasingly large portions of the distance interval are included.

**Figure 10:**
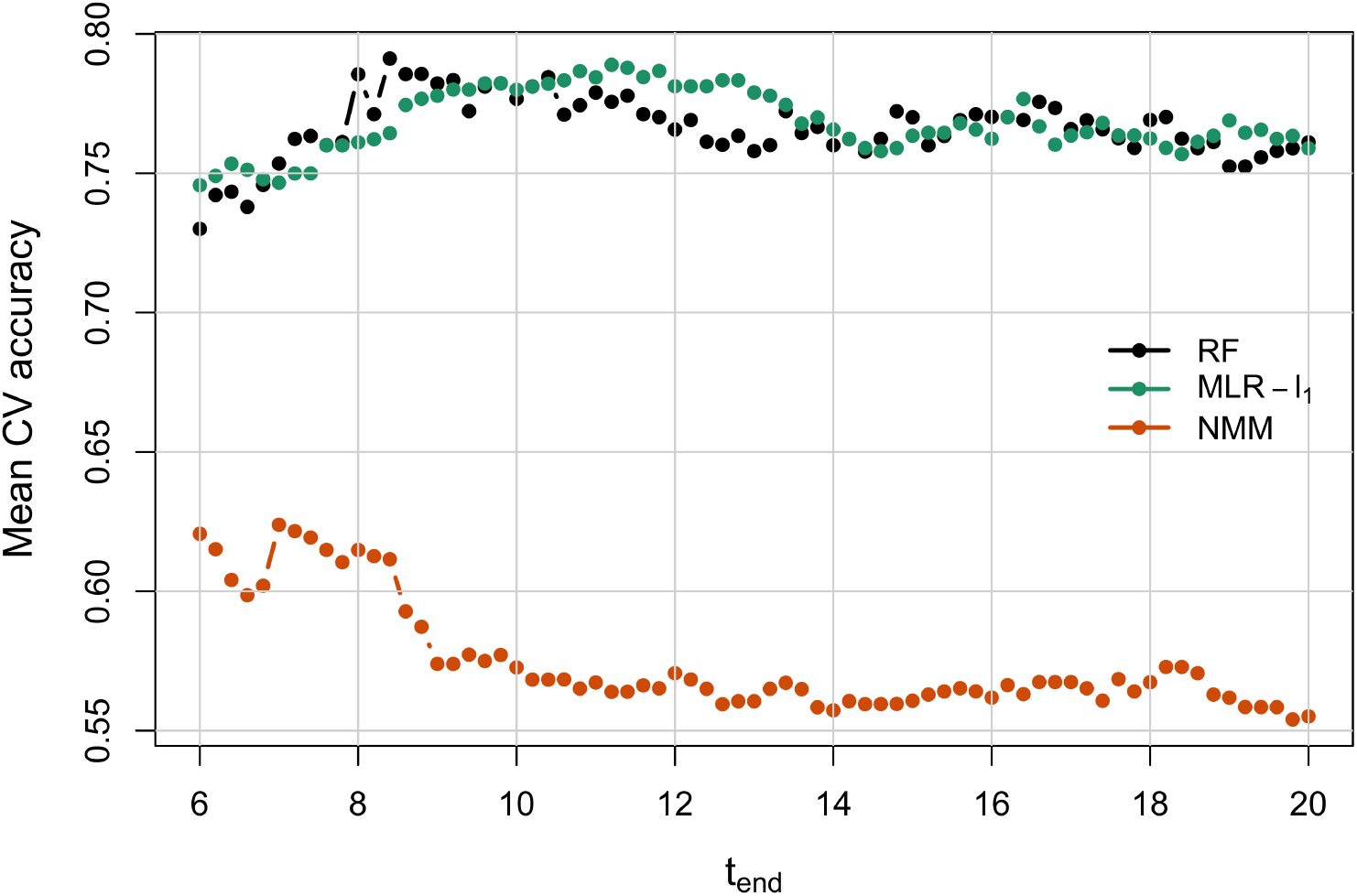
Comparison of five-fold classification accuracy across different *t*_max_ values for the EK dataset using multinomial logistic regression with *l*_1_ penalty (MLR-*l*_1_), nearest Mahalanobis mean (NMM), and random forest classifiers (RF). These results are used for exploratory illustration of the effect of the distance interval and are not the nested cross-validation estimates reported in the subsequent classification analyses.

**Figure 11:**
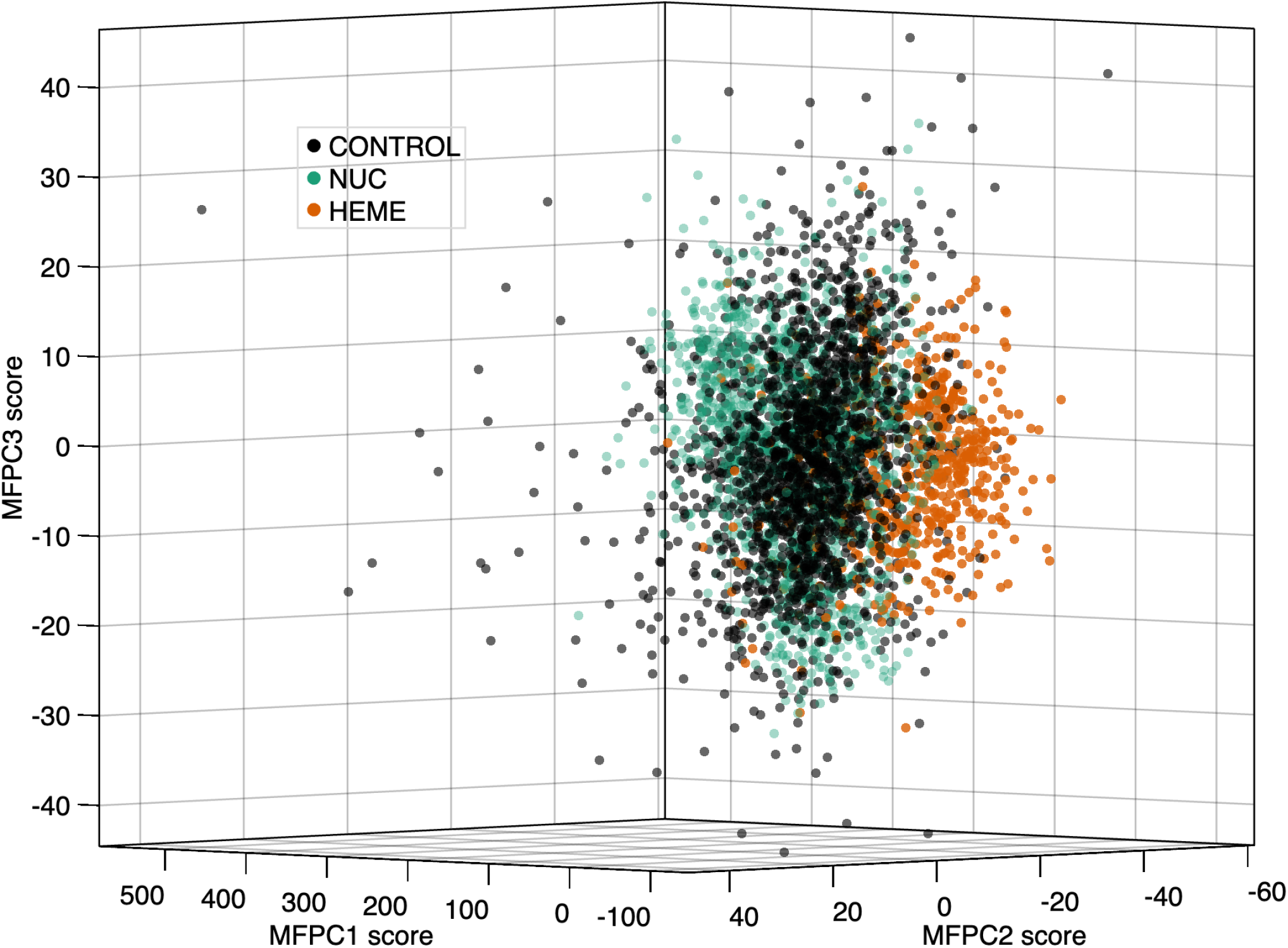
Three-dimensional scatter plot of the first three MFPC scores for the TOUGH-C1 dataset using an illustrative interval 4.0 to 10.0 Å.

Random forest and multinomial logistic regression initially improve as the interval is extended, with performance becoming relatively stable over larger upper endpoints. The nearest Mahalanobis mean classifier exhibits a different pattern, reaching its stronger performance over a shorter interval before declining as the interval is extended further. These differences illustrate that the additional information obtained by extending the FUSED representation need not be equally useful to different classification procedures.

This procedure dependence is also evident when the upper endpoint is selected separately for each classifier. Table 5 summarizes the resulting upper endpoints for the EK and TOUGH-C1 classification tasks. For EK multiclass classification, random forest and multinomial logistic regression select mean upper endpoints of 10.72 Å and 12.16 Å, respectively, whereas the nearest Mahalanobis mean classifier selects a substantially shorter mean endpoint of 7.00 Å. Thus, even for the same dataset and classification task, the portion of the distance interval favored for classification can depend on the classification procedure.

**Table 5:** Selected upper endpoints *t*_max_ for the EK and TOUGH-C1 classification tasks. Values are reported as mean ± standard deviation across the outer folds of repeated nested cross-validation. RF denotes random forest, MLR-*l*_1_ denotes multinomial logistic regression with an *l*_1_ penalty, LR-*l*_1_ denotes binary logistic regression with an *l*_1_ penalty, and NMM denotes the nearest Mahalanobis mean classifier.

| Dataset/task | Classifier | Selected $t_{\max}$ ($\text{\AA}$) |
| --- | --- | --- |
| EK multiclass | RF | $10.72 \pm 2.37$ |
| | MLR-$\ell_1$ | $12.16 \pm 2.82$ |
| | NMM | $7.00 \pm 0.58$ |
| TOUGH-C1 NUC vs Control | RF | $16.32 \pm 3.35$ |
| | LR-$\ell_1$ | $17.52 \pm 1.87$ |
| | NMM | $18.16 \pm 1.99$ |
| TOUGH-C1 HEME vs Control | RF | $13.20 \pm 1.15$ |
| | LR-$\ell_1$ | $11.28 \pm 0.68$ |
| | NMM | $10.80 \pm 2.00$ |

**Table 6:** Performance comparison of *FUSED* with existing approaches for the EK dataset.

| Method | ACC |
| --- | --- |
| *FUSED*-RF | $0.7780 \pm 0.0203$ |
| *FUSED*-MLR-$\ell_1$ | $0.7773 \pm 0.0232$ |
| *FUSED*-NMM | $0.6171 \pm 0.0327$ |
| Sup-CKL [22] | 0.810 |
| CDPA [41] | 0.726 |

The TOUGH-C1 results further demonstrate that the useful distance interval can depend on the classification task itself. For NUC versus Control, all three classifiers select comparatively large mean upper endpoints, ranging from 16.32 to 18.16 Å. For HEME versus Control, the selected endpoints are consistently shorter, ranging from 10.80 to 13.20 Å, with additional differences among classifiers. Hence, changing the classification target within the same dataset can substantially alter which portion of the distance interval is favored by the classification procedure for discrimination.

Taken together, these results illustrate why identification of an informative distance interval should be tied to the particular use of the FUSED representation. The interval is not determined by the dataset alone, nor does classification yield a single interval that is necessarily appropriate across different tasks and procedures. More broadly, other uses of FUSED may place value on different aspects of the threshold-dependent representation and therefore favor different distance intervals. Accordingly, we do not define a single informative interval for each dataset. In the classification analyses that follow, interval selection is instead incorporated into the analysis for each classification setting, with the selection performed using the training data to avoid using held-out observations to determine the interval.

## 8 Comparative Evaluation of the FUSED Representation

Sections 5 and 6 introduced FUSED as a multivariate functional representation of ligand binding sites and illustrated its use for direct functional examination and low-dimensional visualization using the EK dataset. We now evaluate the representation in downstream classification tasks using both the EK and TOUGH-C1 datasets. These datasets provide complementary evaluation settings: EK supports multiclass ligand discrimination, while TOUGH-C1 provides larger binary classification tasks for nucleotide- and heme-binding sites versus controls. As demonstrated in Section 7, the distance interval used for classification can depend on both the task and the classification procedure, so interval selection is incorporated into the evaluation rather than fixed uniformly across analyses. For consistency and simplicity across the classification studies, we use MFPCA as a common dimension-reduction step before applying standard statistical learning methods. The objective is to assess the discriminatory information available in FUSED and compare its resulting performance with existing approaches, rather than to optimize over the full range of possible dimension-reduction and classification procedures.

For the EK and TOUGH-C1 classification analyses, we use repeated nested cross-validation so that selection of the distance interval is incorporated into the evaluation without using the held-out test observations. For each outer training set, the lower endpoint *t*_min_ was fixed at the smallest practical threshold at which the FUSED descriptors could be reliably constructed for nearly all binding sites, while the upper endpoint *t*_max_ is selected using the inner training procedure. For each candidate interval, MFPCA is fit using only the corresponding training data, and the retained MFPC scores are then used to fit the classifier. The selected interval, fitted MFPCA representation, and classifier are subsequently applied to the held-out outer test fold. Thus, the outer-fold performance incorporates interval selection, dimension reduction, and classifier fitting while preserving the independence of the held-out observations used for evaluation.

### 8.1 Extended Kahraman Dataset

For the EK dataset, we fixed the lower endpoint at *t*_min_ = 4.8 Å and selected the upper endpoint *t*_max_ within the repeated nested cross-validation procedure. Candidate values were *t*_max_ ∈ [6, 20] Å in increments of 1 Å. For each candidate interval [4.8, *t*_max_], FUSED descriptors were considered in increments of 0.1 Å. The FUSED representation was reduced using MFPCA within the training data only. The first *p* MFPC scores explaining at least 95% of the total variation were used as predictor variables. The complete procedure was evaluated using 5 repeated 5-fold nested cross-validation, yielding 25 outer test-fold evaluations.

The random forest and multinomial logistic regression results exceed the reported accuracy of the single-threshold CDPA representation and approach the reported performance of Sup-CKL. In particular, FUSED with random forest achieved a mean accuracy of 77.8%, compared with 72.6% reported for CDPA and 81.0% for Sup-CKL. Thus, the FUSED representation supports improved classification performance relative to the reported single-threshold CDPA result while remaining competitive with the alignment-based Sup-CKL approach.

The similarity between the multinomial logistic regression and random forest results is also informative. Although these classifiers differ substantially in flexibility, their comparable performance suggests that the results are due largely to the observed discriminatory structure, rather than the specific linear or nonlinear classification scheme. This is consistent with our use of classification as an evaluation tool for the representation rather than as the central methodological contribution.

The sensitivity analysis in Table 7 examines whether retaining additional MFPCs materially changes classification performance. Increasing the variance-retention threshold from 95% to 98% approximately doubled the mean number of retained MFPC scores, from 6.06 to 12.56. Despite this increase, random forest performance was essentially unchanged, while multinomial logistic regression showed a small decrease that was not statistically significant at the 0.05 level. In contrast, the nearest Mahalanobis mean classifier improved from 57.3% to 60.2%, with the paired sign-flip randomization test indicating a statistically significant difference. Thus, the effect of retaining additional lower-variance MFPCs depends on the classification procedure: the additional components had little effect on the two stronger-performing classifiers but provided useful discriminatory information for the nearest Mahalanobis mean classifier.

**Table 7:** Sensitivity analysis for the proportion of variance retained in MFPCA on the Extended Kahraman dataset at fixed *t*_end_ = 10 Å using 20 repeated 5-fold cross-validation. Accuracies are reported as mean ± standard deviation over 100 fold evaluations. Retaining 95% of the variation used 6.06 ± 0.24 MFPC scores on average, whereas retaining 98% used 12.56 ± 0.50 MFPC scores on average. The *p*-value is from the paired sign-flip randomization test comparing accuracy under 98% and 95% variance retention. Here, RF denotes random forest, MLR-*l*_1_ denotes multinomial logistic regression with *l*_1_ penalty, and NMM denotes the nearest Mahalanobis mean classifier.

| Classifier | Accuracy, 95% | Accuracy, 98% | $p$-value |
| --- | --- | --- | --- |
| RF | $0.7860 \pm 0.0273$ | $0.7871 \pm 0.0274$ | 0.5413 |
| MLR-$\ell_1$ | $0.7793 \pm 0.0217$ | $0.7753 \pm 0.0223$ | 0.0686 |
| NMM | $0.5732 \pm 0.0323$ | $0.6016 \pm 0.0358$ | 0.0001 |

### 8.2 TOUGH-C1 Dataset

We evaluate FUSED on the TOUGH-C1 dataset in two binary classification tasks: nucleotide-binding sites versus control sites and heme-binding sites versus control sites. For each task, we fix the lower endpoint at *t*_min_ = 4.0 Å, the smallest practical threshold at which the FUSED descriptors could be reliably constructed for nearly all TOUGH-C1 binding sites, and select the upper endpoint *t*_max_ within the repeated nested cross-validation procedure. Candidate values were *t*_max_ ∈ [6, 20] Å in increments of 1 Å, with FUSED descriptors evaluated at 0.1 Å increments over each candidate interval [4, *t*_max_]. The complete procedure was evaluated using 5 repeated 5-fold nested cross-validation, yielding 25 outer test-fold evaluations.

Among the three classification procedures, random forest produced the strongest overall performance for both TOUGH-C1 tasks. For nucleotide-binding sites versus controls, random forest achieved a mean outer-fold accuracy of 85.7% and an AUC of 0.935. Performance was stronger for heme-binding sites versus controls, with a mean accuracy of 94.5% and an AUC of 0.978. The corresponding results for multinomial logistic regression and the nearest Mahalanobis mean classifier are reported in Tables 8 and 9. These results show that the FUSED representation supports discrimination in both binary tasks, with particularly strong performance for heme-binding sites.

**Table 8:** Performance comparison of *FUSED* with existing approaches for classifying heme-binding sites in the TOUGH-C1 dataset.

| Algorithm | ACC | PPV | TPR | TNR | AUC |
| --- | --- | --- | --- | --- | --- |
| *FUSED*-RF | $0.9453 \pm 0.0097$ | $0.9485 \pm 0.0249$ | $0.8179 \pm 0.0341$ | $0.9857 \pm 0.0073$ | $0.9781 \pm 0.0080$ |
| *FUSED*-LR-$\ell_1$ | $0.9093 \pm 0.0088$ | $0.8434 \pm 0.0300$ | $0.7673 \pm 0.0207$ | $0.9544 \pm 0.0105$ | $0.9439 \pm 0.0101$ |
| *FUSED*-NMM | $0.8946 \pm 0.0138$ | $0.7427 \pm 0.0336$ | $0.8645 \pm 0.0200$ | $0.9041 \pm 0.0166$ | $0.7984 \pm 0.0419$ |
| BindSiteS-CNN [44] | — | — | — | — | 0.974 |
| BionoiNet [45] | 0.913 | 0.777 | 0.882 | — | 0.960 |
| DeepDrug3D [42] | 0.956 | 0.815 | 0.909 | 0.964 | 0.987 |
| ScanProsite [8] | 0.843 | 0.990 | 0.330 | 0.999 | — |
| G-LoSA [32] | 0.891 | 0.830 | 0.671 | 0.958 | 0.917 |
| Vina [50] | 0.602 | 0.336 | 0.711 | 0.569 | 0.749 |

**Table 9:** Performance comparison of *FUSED* with existing approaches for classifying nucleotide-binding sites in the TOUGH-C1 dataset.

| Algorithm | ACC | PPV | TPR | TNR | AUC |
| --- | --- | --- | --- | --- | --- |
| *FUSED*-RF | $0.8568 \pm 0.0151$ | $0.8432 \pm 0.0186$ | $0.8409 \pm 0.0208$ | $0.8700 \pm 0.0172$ | $0.9346 \pm 0.0099$ |
| *FUSED*-LR-$\ell_1$ | $0.7286 \pm 0.0161$ | $0.7114 \pm 0.0195$ | $0.6764 \pm 0.0270$ | $0.7719 \pm 0.0202$ | $0.7967 \pm 0.0163$ |
| *FUSED*-NMM | $0.7259 \pm 0.0129$ | $0.6833 \pm 0.0137$ | $0.7382 \pm 0.0222$ | $0.7157 \pm 0.0171$ | $0.6222 \pm 0.0241$ |
| BindSiteS-CNN [44] | — | — | — | — | 0.930 |
| BionoiNet [45] | 0.856 | 0.837 | 0.843 | — | 0.935 |
| DeepDrug3D [42] | 0.943 | 0.951 | 0.896 | 0.971 | 0.986 |
| ScanProsite [8] | 0.722 | 0.917 | 0.411 | 0.970 | — |
| G-LoSA [32] | 0.730 | 0.750 | 0.589 | 0.843 | 0.770 |
| Vina [50] | 0.634 | 0.578 | 0.648 | 0.622 | 0.701 |

The random forest results compare favorably with existing approaches reported for TOUGH-C1, achieving performance that is either competitive with or stronger than several established methods. For heme-binding site classification, FUSED-RF achieved an accuracy of 94.5% and an AUC of 0.978, exceeding the reported results for BionoiNet, ScanProsite, G-LoSA, and Vina, while approaching the performance of DeepDrug3D and the reported AUC of BindSiteS-CNN. For nucleotide-binding site classification, FUSED-RF achieved performance essentially identical to BionoiNet, with accuracies of 85.7% and 85.6% and AUC values of 0.935 for both methods after rounding, while exceeding the reported accuracies of BindSiteS-CNN, ScanProsite, G-LoSA, and Vina where accuracy was available. DeepDrug3D achieved higher performance on this task. Overall, these comparisons show that the FUSED representation supports classification performance that is competitive with, and in several cases stronger than, existing approaches while using a standard statistical learning procedure and avoiding pairwise structural alignment.

## 9 TOUGH-M1 pocket matching

We next evaluate FUSED on the TOUGH-M1 benchmark, which provides a substantially different application from the classification tasks considered for the EK and TOUGH-C1 datasets. TOUGH-M1 defines a pairwise pocket-matching problem in which the input is a pair of binding sites (*i, j*) and the objective is to assign a similarity score *s*_*ij*_ such that pockets binding chemically similar ligands receive higher scores than pockets binding chemically dissimilar ligands. The official TOUGH-M1 labels are binary, with *y*_*ij*_ = 1 denoting a positive pair associated with chemically similar ligands and *y*_*ij*_ = 0 denoting a negative pair associated with chemically dissimilar ligands. Performance is therefore evaluated by the ability of the resulting similarity scores to rank positive pocket pairs above negative pocket pairs.

For TOUGH-M1, we fixed the lower endpoint at *t*_min_ = 4.8 Å for construction of the FUSED representation. As in the

EK and TOUGH-C1 analyses, this value was also chosen to balance using a relatively small lower endpoint with retaining as many pockets as possible for which valid FUSED descriptors could be constructed. The upper endpoint was selected within each training split from the candidate set *t*_max_ ∈ {8, 10, 12, 15} Å. For each candidate interval [4.8, *t*_max_], FUSED descriptors were evaluated at increments of 0.1 Å.

### 9.1 Pocket Matching Methods

We first consider a direct pocket-matching approach based on the MFPCA representation of FUSED. For each candidate distance interval, each pocket is represented by its retained MFPC score vector **q**_*i*_. Similarity between pockets *i* and *j* is then defined as the negative Euclidean distance between their MFPC score vectors,

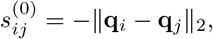

so that larger values of 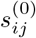 correspond to greater similarity between the two pockets. This provides a direct pocket-matching approach based on the low-dimensional FUSED representation without using the TOUGH-M1 pair labels to train a supervised similarity model.

We next consider whether the full FUSED representation can support supervised learning of pocket similarity using the TOUGH-M1 positive and negative pair labels. For this purpose, we use a Siamese neural network. Siamese neural networks were originally introduced for pairwise verification problems and have since been widely used for learning similarity between paired inputs [4, 17, 37]. This architecture is naturally suited to TOUGH-M1 because the benchmark itself is formulated as a pairwise pocket-matching task.

Our supervised model used a Siamese multilayer perceptron architecture, summarized in Table A2. Each pocket was represented by its raw FUSED feature vector 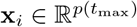, where the input dimension *p*(*t*_max_) is determined by the selected upper endpoint *t*_max_. The two pockets in a pair were passed through the same shared encoder, and the resulting embeddings were compared to estimate the probability that the pair was positive. Model performance was evaluated using ROC-AUC. Full details of the network architecture, training settings, and ROC-AUC calculation are provided in Appendix C.

### 9.2 Evaluation Based on Sequence Clusters

To reduce homologous leakage, we followed the evaluation strategy used in previous TOUGH-M1 neural-network studies, in which protein structures sharing more than 30% sequence identity are assigned to the same data partition during training and testing [44, 47]. Specifically, we mapped TOUGH-M1 protein chains to 30% sequence-identity clusters obtained from the RCSB PDB. Of the 7,524 TOUGH-M1 pockets, 7,494 could be mapped to current RCSB 30% sequence-identity clusters; the remaining 30 pockets could not be mapped to current cluster identifiers and were excluded from the evaluation. After additionally requiring valid FUSED descriptors, 7,447 pockets remained for the cluster-based analysis.

We then performed 20 repeated random train-test splits at the 30% sequence-identity cluster level. In each repetition, 80% of the clusters were assigned to the training partition and the remaining 20% were held out for testing, ensuring that pockets belonging to the same sequence cluster remained within the same partition. The training partition was further divided into model-fitting and validation subsets using the same cluster-based splitting rule. The validation subset was used to select *t*_max_ from {8, 10, 12, 15}Å and, for the supervised Siamese neural network, to perform early stopping. The held-out test partition was not used for model fitting, interval selection, validation, or early stopping and was used only for final performance evaluation. For the direct MFPC-FUSED similarity method, MFPCA was fit using only the model-fitting pockets within each split. The validation and test pockets were then projected into the fitted MFPCA space before pairwise Euclidean distances were computed.

For each split, a pocket pair was assigned to a partition only when both pockets in the pair belonged to sequence clusters assigned to that same partition. Thus, a pair was included in the model-fitting, validation, or test set only when both of its constituent pockets belonged to the corresponding partition. Pairs spanning different partitions, such as one pocket in the training partition and one in the test partition, were excluded from that split.

### 9.3 TOUGH-M1 Pocket Matching Results

The direct MFPC-FUSED similarity method achieved a mean ROC-AUC of 0.7667 ± 0.0234 across 20 repeated random training and test splits based on RCSB 30% sequence-identity clusters. The supervised Siamese FUSED model substantially improved pairwise pocket-matching performance, achieving a mean ROC-AUC of 0.9375 ± 0.0133 across the same 20 held-out evaluations.

The two approaches also favored different portions of the distance domain. Across the 20 repeated splits, the direct MFPC-FUSED similarity method selected a mean upper endpoint of *t*_max_ = 8.45 ± 1.61 Å, whereas the supervised Siamese FUSED model selected *t*_max_ = 10.25 ± 2.49 Å. These values were selected using only the validation partitions within each training split. The larger average upper endpoint selected by the Siamese model suggests that supervised pairwise learning was able to make use of information extending farther from the ligand than was favored by the direct MFPCA-based similarity measure.

These results show that the FUSED representation contains useful information for binding-site matching even without supervised training on the TOUGH-M1 pair labels. At the same time, the substantial improvement obtained by the Siamese model indicates that the full FUSED representation supports considerably stronger pairwise discrimination when the pair labels are used for supervised learning. We further compared the TOUGH-M1 performance of FUSED with previously reported results from established pocket-matching methods (Table 10). Although exact evaluation implementations differ across studies, the supervised Siamese FUSED model achieved performance near the strongest reported results on the benchmark. Its mean ROC-AUC of 0.9375 exceeds the reported values for PocketMatch, APoc, G-LoSA, SiteEngine, Uni-Mol, BindSiteS-CNN, and DeeplyTough, and approaches the reported ROC-AUC of 0.94 for ProFSA. The direct MFPC-FUSED similarity method, despite using no supervised pairwise training, also exceeds several classical pocket-matching approaches.

**Table 10:**
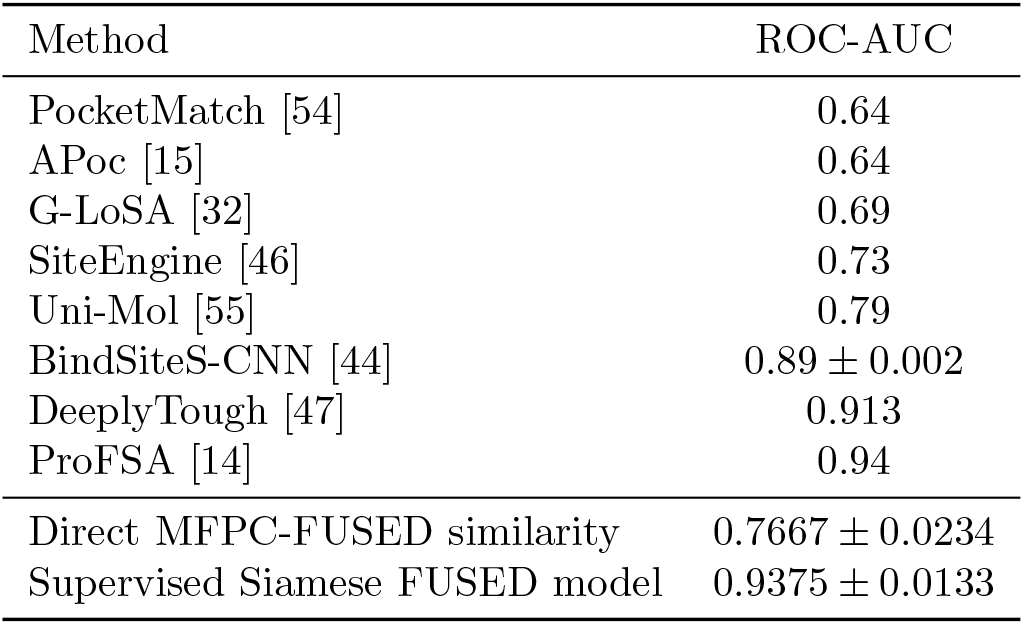
Comparison with published TOUGH-M1 pocket-matching results.

## 10 Computational Considerations

A practical advantage of FUSED is that it produces an object-level representation for each binding site without requiring pairwise structural alignment or pairwise comparison. Once constructed, the resulting representation can be reused across subsequent statistical and machine-learning analyses.

For the EK dataset, the complete five repeated nested cross-validation analysis required 806.4 seconds (13.4 minutes), corresponding to an average of 32.3 seconds per outer test-fold evaluation. Much of this additional computation results from repeatedly fitting the representation and classifier over candidate values of *t*_max_ during interval selection. When the distance interval is specified in advance, the computational cost is substantially lower. For example, construction of the raw FUSED descriptors over the interval 4.8–10.0 Å required 340.4 seconds (5.67 minutes) for the 904 EK binding sites. Twenty repeated 5-fold random-forest evaluations required an additional 310.6 seconds (5.18 minutes), or 3.106 seconds per fold, for a total runtime of 651.0 seconds (10.85 minutes).

For TOUGH-M1, the complete supervised Siamese FUSED analysis over 20 repeated cluster-based train-test splits required approximately 46.01 minutes, corresponding to an average runtime of 2.30 minutes per split.

This computational structure differs substantially from alignment-based approaches such as Sup-CK, for which pairwise binding-site comparisons must be performed explicitly. For 904 binding sites, an exhaustive pairwise analysis requires 408,156 distinct comparisons. Using the reported runtime range of 0.2 to 1.3 seconds per pair, even the lower end of this range corresponds to more than 22 hours of pairwise computation. In contrast, FUSED constructs a representation for each binding site independently, after which the resulting features can be reused with a variety of statistical and machine-learning procedures.

A related distinction arises in comparison with neural-network pocket-matching methods such as DeeplyTough. DeeplyTough reports a runtime of 206.4 seconds for embedding 100 pockets and evaluating 10,000 pairwise comparisons using an NVIDIA Titan X, while training the DeeplyTough model requires approximately one day on a single GPU [47]. Because these timings were obtained using different hardware, datasets, and computational tasks, they should not be interpreted as exact head-to-head runtime comparisons. Nevertheless, they demonstrate a substantial practical computational advantage for FUSED. By explicitly constructing structural and compositional descriptors for each binding site prior to downstream analysis, FUSED avoids the substantial model-training cost associated with deep-learning approaches such as DeeplyTough. This allows FUSED to achieve pocket-matching performance comparable to the strongest reported methods while requiring far less computational time in the analyses considered here.

Overall, these results show that FUSED provides an alignment-free and computationally practical representation that supports competitive downstream performance. Its practical advantage lies not only in the resulting classification and pocket-matching accuracy, but also in the ability to construct reusable object-level descriptors and to examine how useful information changes across the distance-threshold domain.

## 11 Discussion

Ligand binding site representations typically require selecting a fixed distance threshold around the ligand, despite the fact that both local chemical composition and structural geometry evolve as the surrounding neighborhood expands. This creates an inherent tradeoff: thresholds chosen too narrowly may exclude informative structural or compositional information, while broader thresholds may introduce less relevant or redundant information. The proposed FUSED framework addresses this limitation by treating structural and compositional descriptors as functional objects defined over a continuum of distance thresholds, allowing threshold-dependent behavior itself to become part of the representation rather than reducing each binding site to a summary at a single prespecified threshold.

The analyses presented here demonstrate that structural and compositional information exhibit different patterns across the threshold domain. For the EK dataset, the ILR-based compositional descriptors show the most visually pronounced differences among ligand groups near the ligand, with these differences becoming less apparent in the visual summaries as increasingly distant atoms are incorporated. In contrast, the CDPA-based structural descriptors retain more apparent between-group differences over a broader range of thresholds, while also exhibiting threshold-dependent changes in covariance structure that reflect evolving geometric relationships among ligand binding sites. These complementary behaviors support combining the two descriptor families within a unified functional representation. More broadly, they suggest that the information represented by the structural and compositional descriptors changes across distance rather than being fully characterized at a single threshold. FUSED therefore retains not only the structural and compositional descriptors themselves, but also their threshold-dependent behavior as the ligand-centered neighborhood expands.

The analyses across the three benchmark settings instead highlight a broader feature of the framework: informative threshold regions are not expected to be universal, but instead depend on the dataset, analytical objective, and procedure used to analyze the FUSED representation. Across the EK and TOUGH-C1 classification studies, the selected upper endpoints varied with both the classification task and the procedure being applied. The two TOUGH-C1 tasks favored different portions of the distance domain despite being drawn from the same benchmark dataset, while the classifiers used for EK also selected different mean upper endpoints. A similar pattern was observed for TOUGH-M1, where the direct MFPC-FUSED similarity method favored a shorter interval on average than the supervised Siamese FUSED model. Taken together, these results indicate that the useful distance range is determined not by the dataset alone, but by the interaction between the binding-site characteristics represented in the data, the analytical objective, and the procedure used to extract information from the FUSED representation.

The classification studies provide practical validation that the FUSED representation preserves meaningful discriminatory information. Across the EK and TOUGH-C1 benchmark datasets, the representation achieved competitive predictive performance using standard classification procedures without requiring specialized alignment-based comparisons or highly engineered learning architectures. At the same time, the earlier visual analyses demonstrate that predictive accuracy alone does not fully characterize the structure of the representation. Changes in between-group separation or within-group variability may alter the geometry of the feature space in meaningful ways without producing large differences in aggregate classification performance. Thus, the classification results should be viewed as one practical assessment of representation quality, complemented by direct examination of the threshold-dependent functional characteristics of the representation.

The EK and TOUGH-C1 classification studies also illustrate an important consideration in using dimension reduction with FUSED. We used MFPCA as a common unsupervised dimension-reduction procedure for consistency and simplicity across these analyses, but MFPCA identifies directions that explain dominant variation in the functional representation rather than directions selected specifically for discriminatory information. The EK sensitivity analysis illustrates the resulting distinction. Increasing the variance-retention threshold from 95% to 98% approximately doubled the number of retained MFPC scores while producing little change in random forest performance and a small, statistically nonsignificant change in multinomial logistic regression performance. In contrast, the nearest Mahalanobis mean classifier improved significantly when the additional components were retained. Thus, lower-variance components of the FUSED representation can contain useful discriminatory information even when they contribute relatively little to overall functional variation, and the importance of that information can depend on the downstream analytical procedure.

We used TOUGH-M1 to examine the potential of FUSED in a practical pocket-matching setting involving drug-like ligand binding sites. In contrast to the EK and TOUGH-C1 classification tasks, TOUGH-M1 evaluates whether pairs of binding sites are associated with chemically similar ligands. The direct MFPC-FUSED similarity method achieved useful discrimination without supervised training on the TOUGH-M1 pair labels, showing that a low-dimensional unsupervised summary of FUSED retains meaningful information for pocket matching. The supervised Siamese FUSED model, which instead used the full FUSED representation together with the pair labels, substantially increased ROC-AUC and achieved performance near the strongest reported results on the benchmark. Because the two approaches differ both in the representation supplied to the matching procedure and in the use of supervised pairwise learning, the improvement cannot be attributed solely to retaining information omitted by MFPCA. Nevertheless, the results demonstrate that the full FUSED representation can support considerably stronger pairwise discrimination when coupled with a supervised learning procedure. More broadly, they suggest that FUSED may be useful for identifying binding sites associated with chemically similar ligands and for binding-site retrieval applications.

A notable practical advantage of FUSED is that it provides an explicitly defined, alignment-free representation that can be constructed once for each binding site and then used across multiple downstream analyses. Because the representation is built from structural and compositional descriptors rather than learned directly from pairwise labels, its threshold-dependent components can also be examined directly before or alongside subsequent statistical or machine-learning procedures. The computational analyses further demonstrate that this object-level construction can substantially reduce the burden associated with approaches requiring repeated pairwise structural comparisons. Even in the TOUGH-M1 analysis, where FUSED was coupled with supervised neural-network learning, the structural and compositional representation was constructed explicitly before the learning stage rather than requiring the network itself to learn these characteristics from a more primitive structural representation. Together, these features make FUSED computationally practical while retaining flexibility in the choice of downstream analytical procedure.

Comparisons with previously reported benchmark results should nevertheless be interpreted with some caution because the sets of binding sites analyzed are not necessarily identical across studies. Benchmark composition can change over time as structures or database entries used in earlier studies become unavailable, and construction of the FUSED representation also imposes validity requirements that can require additional exclusions. For example, the steroid group in the EK dataset was excluded because too few observations were available for the multiclass analysis, and binding sites for which valid FUSED descriptors could not be constructed over the required distance domain were excluded from the study. Consequently, the comparisons with previously reported methods reflect performance on the same or closely related benchmark tasks rather than exact head-to-head evaluations using identical sets of binding sites.

Several limitations and directions for further work remain. The present implementation of FUSED is based specifically on CDPA-derived structural descriptors and ILR-transformed elemental compositions. The broader functional framework could also incorporate alternative physicochemical, geometric, residue-level, or other threshold-dependent descriptors, and their relative contributions would need to be evaluated for the corresponding analytical tasks. In addition, the distance intervals considered here were selected from prespecified candidate ranges using the corresponding training or validation procedures. Other applications may benefit from alternative strategies for defining or selecting portions of the distance domain. Finally, although the EK, TOUGH-C1, and TOUGH-M1 analyses span multiclass classification, binary classification, and pairwise pocket matching, evaluation on additional datasets and ligand binding-site analysis tasks will be important for determining how broadly the observed advantages generalize.

Overall, FUSED provides a flexible, alignment-free framework for jointly representing structural and compositional characteristics of ligand binding sites across a distance domain rather than reducing each site to a single fixed-threshold summary. The analyses demonstrate that these characteristics change differently across distance, that the portions of the domain most useful for subsequent analysis depend on the dataset, task, and analytical procedure, and that the resulting representation can support both direct functional examination and competitive downstream classification and pocket matching. Together with its computationally practical object-level construction, these results support a broader role for threshold-dependent functional representations in ligand binding-site analysis and chemical informatics.

## Data and Software Availability

All code and metadata necessary to reproduce the analyses in this study, including ligand binding site extraction, descriptor construction, functional representation development, multivariate functional principal component analysis, and classification workflows, are publicly available at https://github.com/tmspriyankara/FUSED-ProteinBindingSite.

The repository includes metadata identifying the benchmark protein structures and ligand binding sites retained for analysis, together with preprocessing inclusion/exclusion information. The original benchmark datasets analyzed in this study are publicly available from their respective public sources, as described in the repository documentation.

## A Classifier Details

Let each binding site *i* = 1, …, *n* belong to one of *K* ligand classes, denoted by *k*_*i*_ ∈ {1, …, *K*}, and let ***ξ***_*i*_ ∈ ℝ^*p*^ be the associated predictor vector.

We use multinomial logistic models as follows

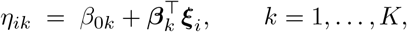

Here, ***β***^*k*^ = (*β*_*k*1_, …, *β*_*kp*_)^⊤^, with *β*_*kj*_ denoting the coefficient associated with predictor *j* in class *k*. Then the class probabilities are:

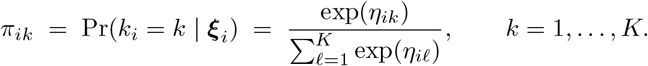

Let *y*_*ik*_ = **1**{*k*_*i*_ = *k*}. The unpenalized multinomial log-likelihood is

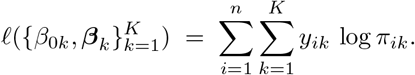

To enable automatic feature selection, we estimate the parameters by maximizing a penalized multinomial log-likelihood with an *l*_1_ penalty [13, 20]

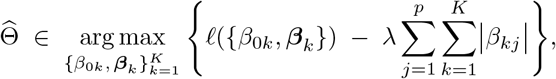

where *λ* ≥ 0 controls overall regularization strength, and it is chosen via cross-validation.

In the nearest Mahalanobis mean classification for each group *k* ∈ {1, …, *K*}, we compute the group-wise mean ***µ***_*k*_ for each class and a common covariance matrix Σ from the training predictor vectors. A new observation ***ξ***_*i*_ ∈ ℝ^*p*^ is assigned to the group minimizing the Mahalanobis distance 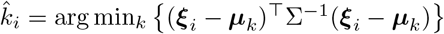

We also use the random forest model, which is a combination of decision trees built on bootstrap samples. Each tree votes for the predicted class, and the final predicted label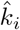 is determined by majority vote [3]. At each split, a random subset of predictors is considered. The number of trees was fixed at 400 and the number of candidate variables considered at each split was set to 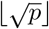. The minimum node size was selected within the inner cross-validation from {2, 5, 7}.

## B Details of Multivariate Functional PCA

In this appendix, we provide the technical details of the multivariate functional principal component analysis (MFPCA) used in Section 6.2. For each ligand binding site, we consider the collection of eight functions consisting of two ILR components and six CDPA components. Let

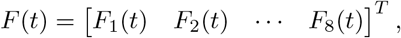

where each *F*_*k*_ ∈ *L*^2^(*T*), and *T* = [0, 1] denotes the rescaled domain of distance thresholds. We view *F* as a random element in the Hilbert space ℋ = [*L*^2^(*T*)]^8^. Let *µ*(*t*) = (*µ* (*t*), …, *µ* (*t*))^*T*^ denote the mean function, where *µ*_k_ (*t*) = E[*F*_k_ (*t*)].

The covariance operator is defined by

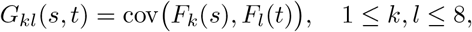

for *s, t* ∈ *T*, yielding a matrix-valued kernel *G*(*s, t*) = *G*_*kl*_(*s, t*) acting on ℋ.

The MFPCA basis functions are obtained as solutions to the integral eigenvalue problem

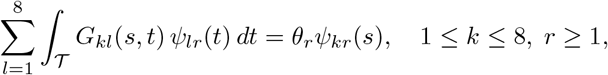

where *Ψ*_*r*_ = (*Ψ*_1*r*_, …, *Ψ*_8*r*_)^*T*^ ∈ ℋ are the multivariate eigenfunctions and *θ*_*r*_ are the corresponding eigenvalues.

The functional representation can then be expressed as

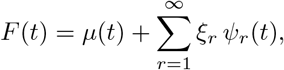

where the MFPC scores *ξ*_*r*_ are given by

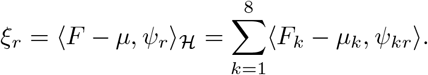

In practice, this expansion is truncated to the first *p* components.

To ensure that different components contribute comparably in the MFPCA, we account for differences in scale between the ILR and CDPA functions. Specifically, we compute average absolute amplitudes across all observations and grid points for the ILR and CDPA components, and define a scaling factor

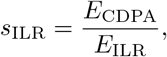

where

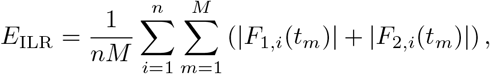

and

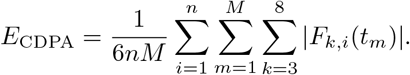

The ILR components are then rescaled by *s*_ILR_, while the CDPA components are left unchanged.

**Table A1:**
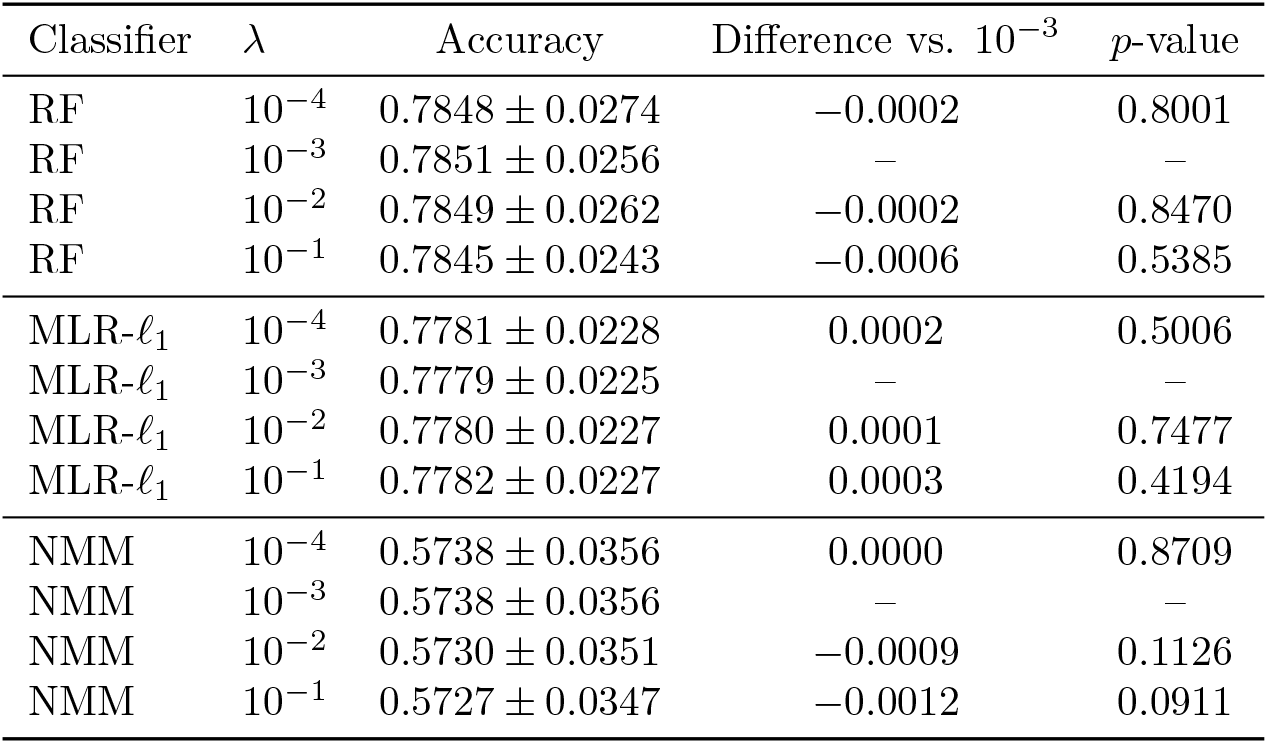
Sensitivity analysis for the B-spline smoothing penalty *λ* on the Extended Kahraman dataset at fixed *t*_end_ = 10 Å using 20 repeated 5-fold cross-validation. Accuracies are reported as mean ± standard deviation over 100 folds. The *p*-value tests the paired accuracy difference relative to the reference value *λ* = 10^*−*3^. Here, RF denotes random forest, MLR-*l*_1_ denotes multinomial logistic regression with *l*_1_ penalty, and NMM denotes the nearest Mahalanobis mean classifier.

| Classifier | $\lambda$ | Accuracy | Difference vs. $10^{-3}$ | $p$-value |
| --- | --- | --- | --- | --- |
| RF | $10^{-4}$ | $0.7848 \pm 0.0274$ | $-0.0002$ | 0.8001 |
| RF | $10^{-3}$ | $0.7851 \pm 0.0256$ | — | — |
| RF | $10^{-2}$ | $0.7849 \pm 0.0262$ | $-0.0002$ | 0.8470 |
| RF | $10^{-1}$ | $0.7845 \pm 0.0243$ | $-0.0006$ | 0.5385 |
| MLR-$\ell_1$ | $10^{-4}$ | $0.7781 \pm 0.0228$ | 0.0002 | 0.5006 |
| MLR-$\ell_1$ | $10^{-3}$ | $0.7779 \pm 0.0225$ | — | — |
| MLR-$\ell_1$ | $10^{-2}$ | $0.7780 \pm 0.0227$ | 0.0001 | 0.7477 |
| MLR-$\ell_1$ | $10^{-1}$ | $0.7782 \pm 0.0227$ | 0.0003 | 0.4194 |
| NMM | $10^{-4}$ | $0.5738 \pm 0.0356$ | 0.0000 | 0.8709 |
| NMM | $10^{-3}$ | $0.5738 \pm 0.0356$ | — | — |
| NMM | $10^{-2}$ | $0.5730 \pm 0.0351$ | $-0.0009$ | 0.1126 |
| NMM | $10^{-1}$ | $0.5727 \pm 0.0347$ | $-0.0012$ | 0.0911 |

## C Pocket Matching Methods for TOUGH-M1 data set

### C.1 Direct MFPC-FUSED Similarity

In this method, we represented each pocket by its FUSED descriptors and projected onto the retained MFPC score space. Let *q*_*i*_ be a resulting score vector of the *i*^*th*^ pocket. Similarity between a pair of pockets *i* and *j* was then defined as the negative Euclidean distance between their MFPCA scores,

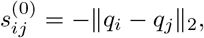

with larger values of 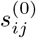 indicating greater pocket similarity.

### C.2 Supervised Siamese Neural Network with FUSED

Here, we used a Siamese multilayer perceptron architecture, which is summarized in Table A2. Each pocket was represented by its raw FUSED feature vector 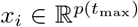, where *p*(*t*_max_) is determined by the selected upper endpoint *t*_max_. The two pockets in a pair were passed through the same shared encoder *g*(·), producing 64-dimensional embeddings,

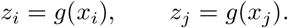

The encoder consisted of three fully connected layers with ReLU activations, batch normalization, and dropout. The two embeddings were then compared using three pairwise features absolute difference (|*z*_*i*_ −*z*_*j*_|), elementwise product (*z*_*i*_ ⊙*z*_*j*_), and cosine similarity (cos(*z*_*i*_, *z*_*j*_)). These features are commonly used to compare paired embeddings in neural networks [19,24,43]. These three comparison features were concatenated into a 129-dimensional pairwise feature vector,

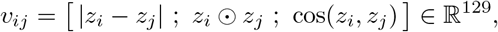

and passed to a binary classification head to estimate the pairwise matching score

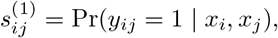

the predicted probability that pockets *i* and *j* form a positive pair.

The network was trained using binary cross-entropy loss,

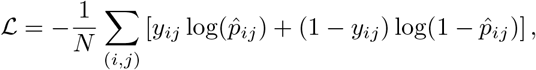

where 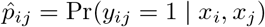 is the predicted probability that the pair is positive and *y*_*ij*_ 0, 1 is the official TOUGH-M1 pair label. Optimization used AdamW [36] with learning rate 10^*−*3^, weight decay 10^*−*4^, dropout rate 0.20, batch size 4096, and a maximum of 20 epochs, with early stopping after 4 consecutive epochs without validation improvement.

## C.3 ROC-AUC Calculation

Performance was evaluated using the area under the receiver operating characteristic curve (ROC-AUC). For each method, every evaluated pocket pair (*i, j*) received a scalar score *s*_*ij*_. Larger values of *s*_*ij*_ indicate stronger evidence that the pair was positive. For the direct MFPCA-FUSED baseline, this score was the negative Euclidean distance 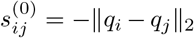. For the supervised Siamese neural network, the score was the predicted positive-pair probability 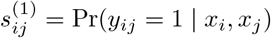.

ROC-AUC was then computed by ranking all evaluated positive and negative pocket pairs according to *s*_*ij*_. ROC-AUC estimates the probability that a randomly selected positive pair receives a higher score than a randomly selected negative pair. Let

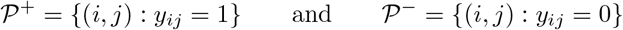

denote the evaluated positive and negative pair sets, where *n*^+^ is the number of elements in the set *P* ^+^ and *n*^*−*^ is the number of elements in the set *P* ^*−*^. The empirical ROC-AUC was computed as the Wilcoxon statistic as follows. [18]

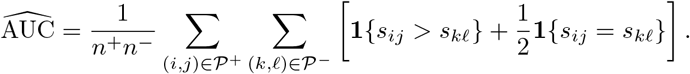

An AUC of 0.5 corresponds to random ranking, whereas an AUC of 1.0 indicates that all evaluated positive pairs are ranked above all evaluated negative pairs.

**Table A2:**
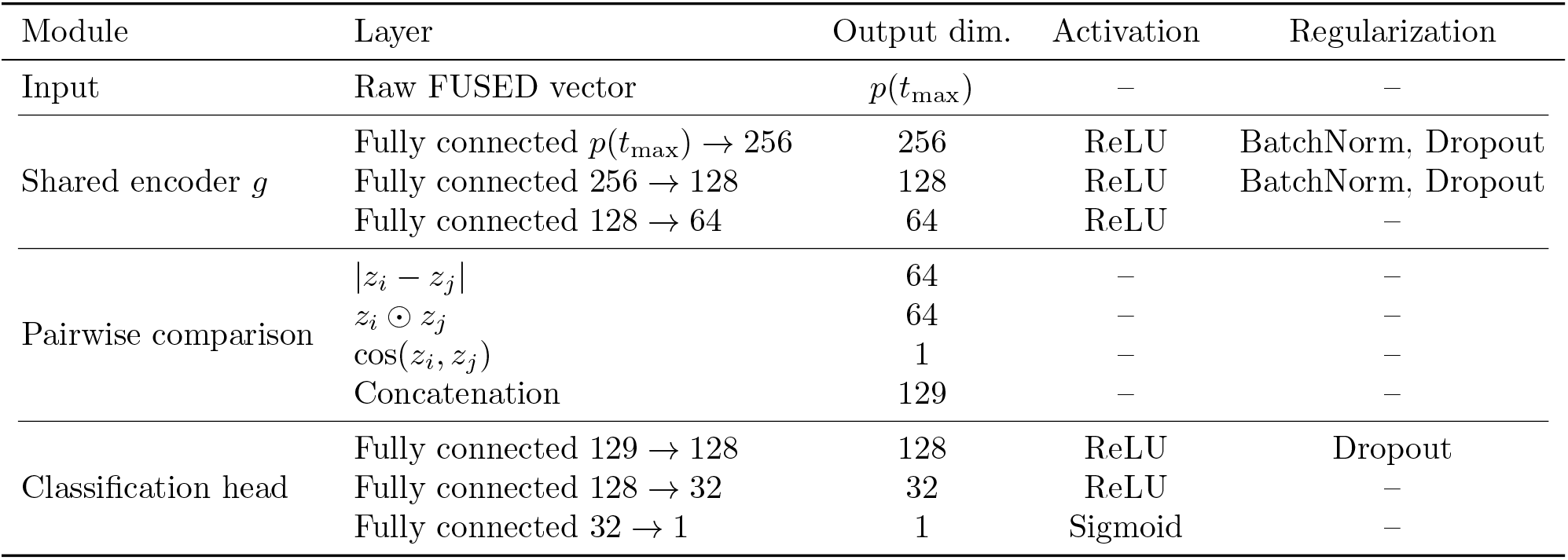
Siamese FUSED architecture for TOUGH-M1 pairwise pocket matching. Each pocket is represented by a raw FUSED vector of dimension *p*(*t*_max_). The raw FUSED input dimension is *p*(*t*_max_) = 8|T (*t*_max_)|, where T (*t*_max_) is the 0.1 Å threshold grid over [4.8, *t*_max_]. The shared encoder *g* maps each pocket to a 64-dimensional embedding. The two embeddings are compared using absolute difference, elementwise product, and cosine similarity, then concatenated into a 129-dimensional pairwise feature vector and passed to the classification head.

## Notes

### Competing Interest Statement

The authors have declared no competing interest.

### Summary of Updates

This revised version includes an expanded evaluation of FUSED on the TOUGH-M1 pocket-matching benchmark, including a comparison with published pocket-matching methods. The cross-validation procedures were revised to use repeated and nested cross-validation where appropriate. Additional sensitivity analyses were added for MFPCA variance retention and B-spline smoothing. The Methods section was revised to clarify the general FUSED functional representation and computational workflow. Results tables, figures, and runtime analysis were updated accordingly.

